# The Z-shaped N-terminal Domain of Atg11 Coordinates Atg9 Binding and Recruitment in Selective Autophagy

**DOI:** 10.64898/2026.08.17.744853

**Authors:** Sarah I Najera, Devika Andhare, Anna E. Hill, Zhanibek Bekkhozhin, Michael J Ragusa

## Abstract

Macroautophagy is a conserved catabolic process that facilitates the degradation of cellular material by capturing it in double membrane vesicles termed autophagosomes. In *Saccharomyces cerevisiae*, selective macroautophagy is initiated by the scaffolding protein Atg11. Atg11 recruits the transmembrane protein Atg9, which resides in small vesicles, to autophagic cargo. Atg9 vesicles then fuse, forming the initial membrane sheet that expands into the autophagosomal membrane. While it is known that Atg9 interacts with Atg11 via a set of hydrophobic amino acids in the disordered N-terminus of Atg9, it is unclear how Atg11 mediates this interaction. To gain insight into this unknown aspect of autophagy initiation we utilized a combination of biochemical, structural, and cellular approaches. We demonstrate that the N-terminal domain (NTD) of Atg11 is the primary interaction site for Atg9, but the NTD requires clustering by the C-terminal region of Atg11 for its complete interaction with Atg9. We investigated the structure of the Atg11-NTD using cryo-EM which, in combination with AlphaFold modeling, revealed a positively charged binding pocket within the Atg11-NTD that is essential for Atg9 binding. Mutation of this conserved binding pocket leads to a loss of Atg9 binding in yeast and a reduction in the selective autophagy of mitochondria. Taken together, our results demonstrate the mechanism by which Atg11 recruits Atg9 to autophagy initiation sites.

## INTRODUCTION

Autophagy is the cellular process of “self-eating”, in which cytoplasmic material is targeted to lysosomes for degradation. Macroautophagy (hereafter, autophagy), which is conserved from yeast to humans, is distinct from other forms of autophagy in that cargo to be degraded is captured in double-membered vesicles, termed autophagosomes [1,2]. Completed autophagosomes fuse with the vacuole, in yeast, or lysosomes, in higher eukaryotes, to degrade the captured material into free amino acids, fatty acids and other metabolites to be reused by the cell [2,3]. Autophagy is therefore critical to maintaining cellular homeostasis in addition to promoting cellular survival in nutrient deprived environments [4,5]. Unsurprisingly, autophagy defects have been implicated in many human diseases, including neurodegeneration, cancer, and autoimmune diseases [6–8].

Autophagy can degrade a variety of cargos, ranging from damaged or excess mitochondrial fragments, endoplasmic reticulum, nuclear fragments, and other cytosolic material [9]. Cargo can be captured via non-selective autophagy, where cargo is captured at random, or selective autophagy, where cargo is marked for degradation by selective autophagy receptor (SAR) proteins [9]. SARs recruit autophagy proteins directly to cargo leading to an autophagosome that is built around the cargo to be degraded. In contrast, non-selective autophagy, which is typically induced by nutrient limiting conditions, results in the capture of material without direct recognition of the cargo.

Autophagosome biogenesis is facilitated by a set of proteins termed the core autophagy machinery [10]. These core autophagy proteins recruit the initial membrane source for autophagosome biogenesis and facilitate the expansion of this membrane into a double membrane sheet. In the yeast *Saccharomyces cerevisiae*, the core autophagy factors are recruited to autophagy initiation sites by one of two autophagy scaffolding proteins: Atg11 or Atg17 [11,12]. In non-selective autophagy, Atg17 recruits the core autophagy machinery to a site adjacent to the vacuole, termed the phagophore assembly site (PAS) [13]. This process leads to an autophagosome that is built in close proximity to the vacuole. In selective autophagy, Atg11 recruits the core autophagy machinery to the cargo to be degraded through its direct interaction with cargo specific SARs [9,14,15]. This recruitment leads to a direct linkage between the cargo and the autophagosomal membrane which is maintained throughout by other core autophagy factors. In mammalian cells, FIP200 appears to serve a dual role, functioning as the scaffolding protein for both non-selective and selective autophagy and helping recruit the mammalian homologs of the core autophagy proteins to cargo [16–18].

Atg11 is a 1178 amino acid protein that consists of four predicted domains, including an N-terminal domain (NTD), a large coiled-coil domain (CC3), a helical region of unknown function, and the C-terminal CLAW domain that binds SARs (**Figure 1A**) [11]. The overall domain architecture of Atg11 is similar to that of FIP200, suggesting that FIP200 is structurally more like Atg11 than Atg17, despite FIP200’s dual role in mammalian cells. Currently, the only experimental structure of Atg11 is the crystal structure of a fragment of the CC3 [19]. However, AlphaFold 3 structure predictions for different domains of Atg11 are similar to the experimentally determined structures of FIP200, further supporting that FIP200 is much more similar to Atg11 than Atg17 [17,20]. Atg11 can be recombinantly expressed in *E. coli* as two stable soluble folded fragments, the Atg11-NTD (1-646) and the C-terminal region (CTR, 699-1178), which contains the CC3 through the end of the protein. Biophysical studies of these constructs demonstrated that each region independently forms parallel dimers in solution, demonstrating that Atg11 contains multiple independent dimerization regions [21].

**Figure 1.**
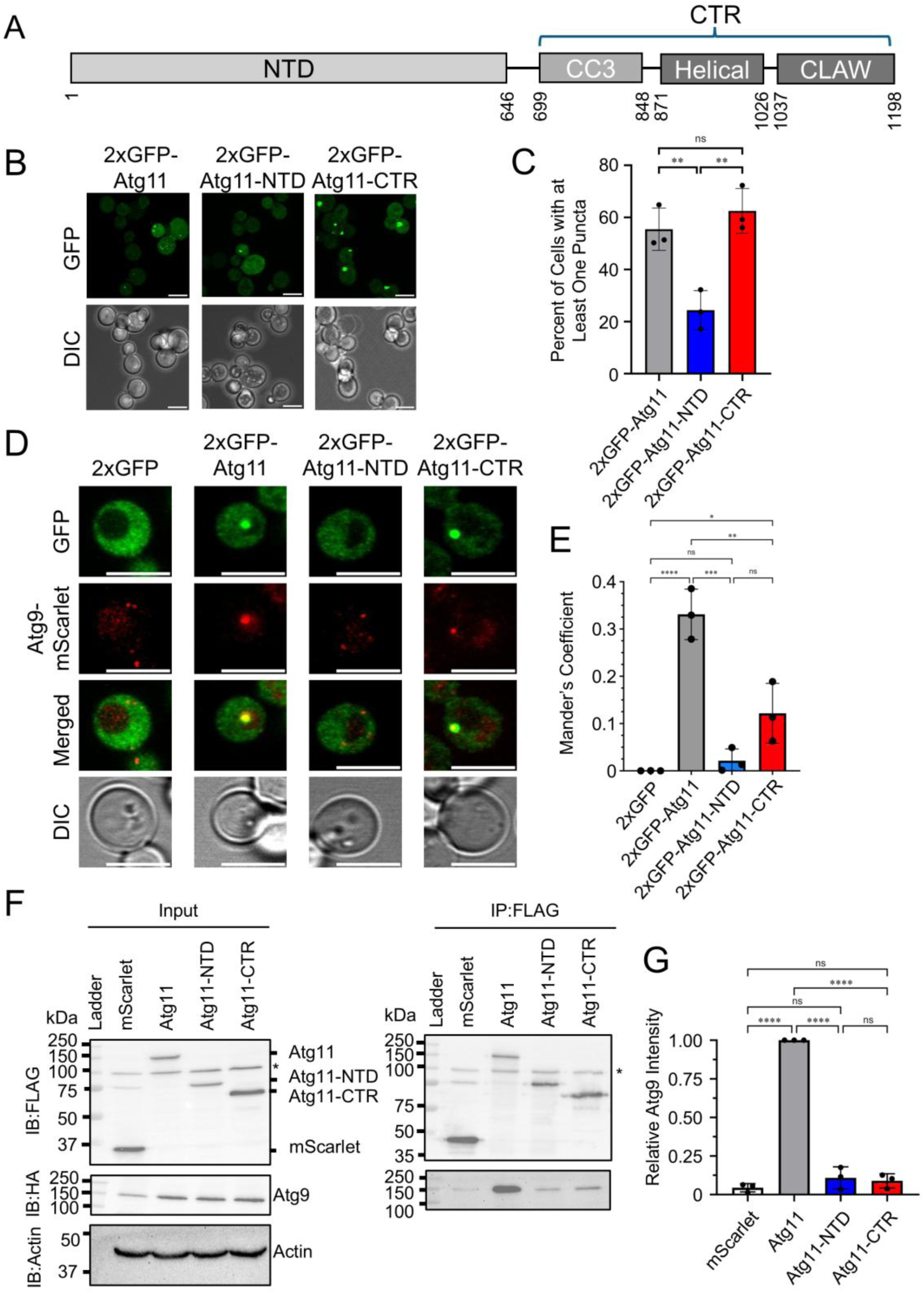
Atg9 colocalizes and interacts with Atg11 when both the NTD and CTR are present. (A) Domain map of Atg11 showing the Atg11-NTD and Atg11-CTR containing CC3, Helical and CLAW domains. The starting and ending amino acid for each domain is written below. (B) Representative microscopy images of *atg11*Δ *S. cerevisiae* cells transformed with the indicated 2xGFP-Atg11 constructs. Scale bars: 5 μm. (C) Quantitation of the images from B displaying the percentage of cells with at least one 2xGFP-Atg11 puncta in nutrient rich (SMD) media. Data represents means ± SD from three independent repeats (n=3). ** indicates p ≤ 0.01, ns indicates non-significant from a One-way ANOVA with Tukey’s multiple comparisons test. (D) Microscopy images of *atg11*Δ *atg9*Δ *S. cerevisiae* cells transformed with the indicated 2xGFP-Atg11 construct and Atg9-mScarlet grown in SMD. Scale bars: 5 μm (E) Mander’s Coefficient quantification of images from D to display co-occurrence of Atg9-mScarlet with 2xGFP-Atg11 constructs. Data represents means ± SD from three independent repeats (n=3). * indicates p ≤ 0.05, ** indicates p ≤ 0.01, *** indicates p ≤ 0.001, **** indicates p < 0.0001, ns indicates non-significant from a One-way ANOVA with Tukey’s multiple comparisons test. (F) Representative Western blots of co-immunoprecipitation assay of *atg11*Δ *atg9*Δ *atg17*Δ *S. cerevisiae* cells transformed with 3xFLAG-mScarlet or the indicated 3xFLAG-Atg11 constructs, and Atg9-HA grown in SMD. Each immunoblot (IB) is labeled with the antibody that was used. Actin was used as a loading control. * indicates non-specific bands. All other bands are labeled. (G) Quantification of the relative Atg9 band intensity from the western blots from F. Data represents means ± SD from three independent repeats (n=3). **** indicates p < 0.0001, ns indicates non-significant calculated using One-way ANOVA with Tukey’s multiple comparisons test.

Atg9 is the only essential transmembrane protein for both selective and non-selective autophagy in yeast and it plays a critical role in both the initiation and elongation of autophagosomes [22,23]. The mammalian homologs of Atg9, ATG9A and ATG9B, serve a similar function in mammalian autophagy [24–27]. In *S. cerevisiae*, Atg9 is embedded in 30-60 nm Golgi-derived vesicles that function as the initial membrane seed during autophagy initiation [22,28,29]. Clusters of Atg9 vesicles can be found throughout the cytosol and upon autophagy induction a subset of these vesicles are recruited to cargo by Atg11 [23,30]. During autophagosome expansion, Atg9 functions as a scramblase that helps equilibrate lipids across the autophagosomal lipid bilayer after they are transported from the ER by the lipid transporter Atg2 [25–27].

Atg11 interacts directly with Atg9 to recruit Atg9 vesicles to selective autophagy initiation sites. This interaction requires two Proline-Leucine-Phenylalanine (PLF) motifs in the N-terminal disordered region of Atg9 [31]. However, it is still unknown which region of Atg11 is responsible for binding Atg9, including how the PLF motifs are directly recognized by Atg11. Therefore, we set out to determine which region of Atg11 interacts with Atg9, how this interaction is coordinated by Atg11, and the impact of disrupting this interaction on selective autophagy in yeast.

## RESULTS

### Both the NTD and CTR of Atg11 are required for the interaction of Atg11 and Atg9 in cells

Atg11 can be expressed in and purified from *E. coli* as two different fragments, the NTD (1-646) and the CTR (699-1178), both of which are soluble, folded and stable dimers (**Figure 1A**) [21,32]. We took advantage of these stable constructs to investigate which of these regions of Atg11 is responsible for binding to Atg9 in cells. Atg11 forms puncta during autophagy initiation and Atg9 colocalizes with these puncta. We previously reported that Atg11 and the Atg11-CTR form puncta in a multiple knockout (MKO) cell line that lacks most autophagy proteins, but the Atg11-NTD does not [32,33]. However, we did not test which of these constructs form puncta when other autophagy proteins are present, and this would be required to monitor the colocalization of Atg11 and Atg9. Therefore, we first monitored different 2xGFP-Atg11 constructs expressed from centromeric plasmids under the control of their endogenous promoter in *atg11Δ S. cerevisiae* (**Figure 1B**). 2xGFP-Atg11 and 2xGFP-Atg11-CTR both formed puncta in *atg11Δ* cells similarly to MKO cells. The percent of cells containing at least one Atg11 puncta was similar between these two constructs, with 55.5 ± 8.1% and 62.5 ± 8.6% of cells having at least one punctum for 2xGFP-Atg11 and 2xGFP-Atg11-CTR, respectively (**Figure 1C**). In contrast, only 24.3 ± 7.6% of cells expressing 2xGFP-Atg11-NTD had at least one punctum (**Figure 1C**). These results demonstrate that 2xGFP-Atg11-NTD forms puncta when other autophagy proteins are present but that the number of puncta is significantly reduced compared to Atg11 and Atg11-CTR. Therefore, each of these Atg11 constructs can be used to examine Atg11 and Atg9 colocalization.

We next tested whether Atg9-mScarlet colocalizes with different 2xGFP-Atg11 puncta. As expected, 2xGFP-Atg11 and Atg9-mScarlet colocalized in cells and had a co-occurrence coefficient (Mander’s coefficient) of 0.33 ± 0.05, where a value of 0 indicates no-overlap and a value of 1 indicates complete signal overlap (**Figure 1D and E**). In contrast, 2xGFP-Atg11-NTD showed no colocalization with Atg9-mScarlet, with a Mander’s coefficient of 0.02 ± 0.02, despite approximately one quarter of these cells containing 2xGFP-Atg11-NTD puncta. 2xGFP-Atg11-CTR showed some colocalization with Atg9-mScarlet with a Mander’s coefficient of 0.12 ± 0.06. The colocalization of 2xGFP-Atg11-CTR was significantly reduced when compared to 2xGFP-Atg11, suggesting that 2xGFP-Atg11-CTR can recruit Atg9 to autophagy initiation sites but that this recruitment is reduced compared to full-length Atg11.

To further explore which region of Atg11 interacts with Atg9 in cells, we performed a co-immunoprecipitation experiment with 3xFLAG-Atg11, 3xFLAG-Atg11-NTD or 3xFLAG-Atg11-CTR and Atg9-HA. 3xFLAG-mScarlet was used as a negative control. Co-immunoprecipitation experiments were performed in *atg11Δ atg9Δ atg17Δ S. cerevisiae* cells to prevent Atg9 turnover via non-selective autophagy, as previously observed [34,35]. Under nutrient rich conditions, Atg9 co-immunoprecipitated with Atg11, as previously reported (**Figure 1F and G**) [31]. However, Atg9 co-immunoprecipitation with either Atg11-NTD or Atg11-CTR was close to the background binding observed when mScarlet was immunoprecipitated (**Figure 1F and G**). This is in good agreement with our colocalization studies and suggests that both the Atg11-NTD and Atg11-CTR are required for the complete interaction between Atg11 and Atg9 in cells.

### Atg11-NTD binds directly to Atg9(1-255) in-vitro

We demonstrated that Atg9 colocalization and co-immunoprecipitation with Atg11 requires both the NTD and CTR. Therefore, to better understand why both the NTD and CTR might be required for interacting with Atg9 in cells, we tested whether Atg9(2-255) directly interacts with the Atg11-NTD and Atg11-CTR using purified proteins. Atg9(2-255) was used, as it was previously shown to contain the primary Atg11 binding region [31]. GFP, GFP-Atg11-NTD, GFP-Atg11-CTR and mScarlet-Atg9(2-255) were recombinantly expressed and purified using *E. coli*. GFP-trap beads were saturated with each of the different GFP tagged constructs, then incubated with mScarlet-Atg9(2-255) and imaged (**Figure 2A**). mScarlet-Atg9(2-255) bound to GFP-Atg11-NTD on GFP-trap beads, but not to GFP or GFP-Atg11-CTR, suggesting that the main interaction region for Atg9 is the Atg11-NTD (**Figure 2B**). To verify the interaction between Atg9(2-255) and Atg11-NTD, we purified GST-Atg9(2-255), labeled it with Alexa-Fluor-594, and repeated the GFP trap assay (**Figure 2C**). We observed similar results with Alexa-Fluor-594 labeled Atg9(2-255), confirming that Atg9(2-255) binds directly to the Atg11-NTD (**Figure 2C and D**).

**Figure 2.**
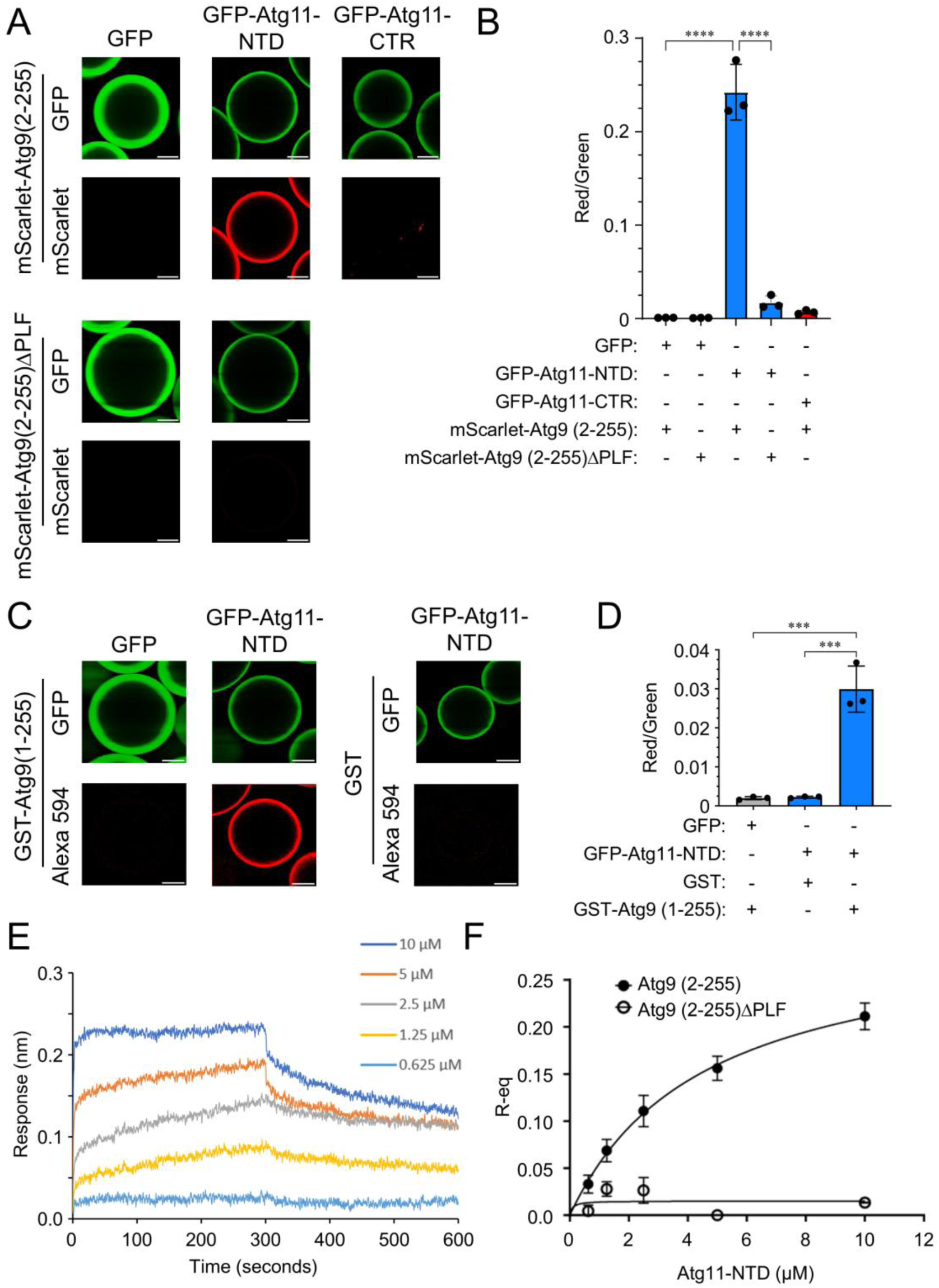
Atg9(2-255) binds directly to the Atg11-NTD. (A) Representative fluorescence microscopy images of GFP-trap beads with the indicated GFP-Atg11 constructs or GFP immobilized then loaded with mScarlet-Atg9(2-255) or mScarlet-Atg9(2-255ΔPLF). Fluorescence intensities were normalized to GFP-Atg11-NTD with mScarlet-Atg9(2-255). Scale bars: 25 µm (B) Quantification of images from A in which fluorescence intensity around the bead was quantified for green and red channels of 5 beads per image with 3 images per repeat. Data represents means ± SD from three independent repeats (*n*= 3). **** indicates p < 0.0001, calculated using One-way ANOVA with Dunnett multiple comparisons test. (C) Fluorescence microscopy images of GFP-trap beads with GFP or GFP-Atg11-NTD immobilized then loaded with Alexa 594 labeled GST-Atg9(1-255) or Alexa 594 labeled GST. Fluorescence intensities were normalized to GFP-Atg11-NTD with GST-Atg9(1-255). Scale bars: 25 µm (D) Quantification of images from C in which fluorescence intensity around the bead was quantified for both the green and red channels of 5 beads per image with 3 images per repeat. Data represents means ± SD from three independent repeats (*n*= 3). *** indicates p ≤ 0.001, calculated using One-way ANOVA with Dunnett multiple comparisons test. (E) Biolayer Interferometry traces of 6xHis-Atg9(2-255) immobilized on Sartorius HIS1K tips then dipped into varying concentrations of Atg11-NTD. (F) Equilibrium (R-eq) of traces from E plotted against Atg11-NTD concentration and fitted using a non-linear regression model resulting in a K_d_ of 4.63 ± 0.757 µM for Atg11-NTD and Atg9(2-255). No K_d_ was able to be determined for Atg11-NTD and Atg9(2-255ΔPLF). Data represents means ± SD from three independent repeats (n=3) for Atg11-NTD and Atg9(2-255) and two independent repeats (n=2) for Atg11-NTD and Atg9(2-255ΔPLF).

We wanted to determine if the interaction between Atg11-NTD and Atg9(2-255) is mediated by the previously identified PLF motifs in Atg9 that were shown to be required for the interaction between Atg9 and Atg11 [31]. Therefore, we deleted both of these motifs (Δ163-165 Δ187-189) from our mScarlet-Atg9(2-255) construct and repeated the GFP trap assay. Deletion of the PLF motifs from Atg9(2-255) led to a complete loss in binding to GFP-Atg11-NTD, demonstrating that the primary interaction region for Atg9 is the Atg11-NTD and that this interaction is mediated by the PLF motifs in Atg9 (**Figure 2A and B**).

Next, we wanted to determine the binding affinity of Atg11-NTD for Atg9(2-255). We performed Biolayer Interferometry (BLI), in which 6xHis-Atg9(2-255) was immobilized on HIS1K Sartorius tips at 10 µM, then dipped in Atg11-NTD at concentrations ranging from 0.625 µM to 10 µM (**Figure 2E)**. Fitting of the steady state data led to a dissociation constant (K_d_) of 4.63 ± 0.757 µM (**Figure 2F**). BLI was also used to further validate the importance of the Atg9 PLF motifs by repeating our BLI experiment with the same conditions, except using 6xHis-Atg9(2-255)ΔPLF instead of 6xHis-Atg9(2-255). 6xHis-Atg9(2-255ΔPLF) had no observable binding with Atg11-NTD, confirming that the PLF motifs mediate the interaction between Atg11-NTD and Atg9 (**Figure 2F and S1**).

### The Atg11-NTD is a Z-shaped dimer

To better understand how the Atg11-NTD interacts with Atg9, we examined the structure of the Atg11-NTD using cryo-electron microscopy. 2D class averages revealed that that the Atg11-NTD is an elongated Z shape, consistent with the AlphaFold model of the Atg11-NTD dimer (**Figure 3A and S2**). After several iterative rounds of 2D classification we were able to identify orthogonal views of Atg11-NTD that were required to obtain an interpretable reconstruction. Following particle cleaning in 2D classification, multiclass ab-initio, heterogenous refinement and non-uniform refinement we were able to obtain a 6.68 Å reconstruction of the Atg11-NTD dimer (**Figure 3B and S3 and Table S1**). The reconstructed map revealed a twisted Z structure with one arm of the Z bent relative to the other. One arm of the Z contained weaker and more broad density than the other, suggesting that the arms of the Z may be mobile relative to the center portion of the Z. The Atg11-NTD AlphaFold 3 model docked well into the reconstruction, with one monomer and the dimerization region fitting well into the density **(Figure 3C)**. The remaining portion of the second monomer is shifted relative to the density, suggesting a slight difference in the overall shape of the Z compared to the prediction **(Figure 3C)**. Given how weak the density of the second monomer is, we performed local refinement focusing on one arm and the central region of the Z **(Figure S3)**. The resulting reconstruction was 6.39 Å and had an improved map. The reconstruction had clear long tubular densities consistent with the size and shape of α-helices (**Figure 3D**). After docking the AlphaFold 3 model of the Atg11-NTD dimer into our local refinement reconstruction we observed density for the majority of one monomer of Atg11-NTD except residues 1-19, 102-149 and 597-646 suggesting that these regions are likely dynamic or heterogenous. This is in good agreement with the low confidence scores for the majority of these regions (**Figure S2**). The region spanning amino acids 507 to 533 had low confidence scores in the Atg11-NTD AlphaFold 3 prediction. However, we see clear density for this region indicating that this region is correctly modeled in the AlphaFold 3 prediction. We also observed density within the dimerization region for 456-596 from the other monomer of the Atg11-NTD, demonstrating that dimerization is supported by amino acids 456-596 from both monomers. To generate a structural model based on our local refinement we used the AlphaFold 3 model and deleted the regions of the structure that had no density. We then performed chain refinement with tight geometrical constraints to improve the agreement between the model and the map without distorting the overall structure. The density for amino acids 535 through 546 was much weaker than the rest of the structure suggesting possible heterogeneity or dynamics in this region. However, we decided to leave this region in the structure because clear density was visible on either side of this region. This resulted in a hybrid structural model based on AlphaFold modeling and our cryo-EM volume which includes one monomer of Atg11-NTD along with the dimerization region (456-596) of the other monomer **(Figure 3D and E)**.

**Figure 3.**
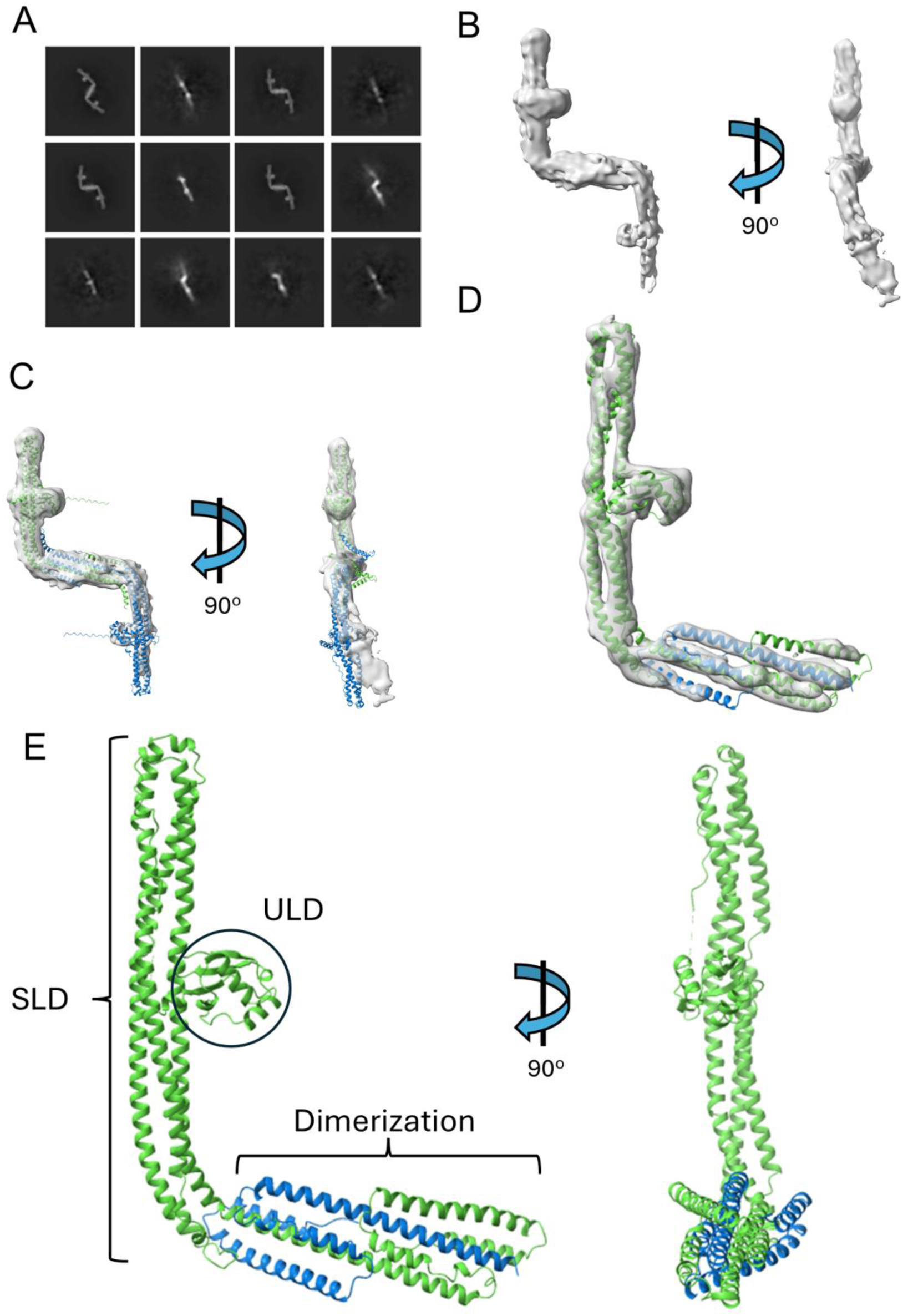
Cryo-EM analysis of the Atg11-NTD. (A) Representative 2D class averages of Atg11-NTD showing both side and orthogonal views. (B) 6.68 Å reconstruction of the Atg11-NTD shown at two different angles. (C) Overlay of the reconstruction from B with the Atg11-NTD dimer AlphaFold 3 prediction shown as a cartoon representation. One monomer of Atg11-NTD is shown in green and the other in blue. (D) 6.39 Å reconstruction from local refinement focusing on one monomer and the dimerization region of the Atg11-NTD. The hybrid structural model of the Atg11-NTD was generated using the AlphaFold model of the Atg11-NTD dimer and regions were removed that were not visible in the density. The Atg11-NTD hybrid model is shown as a cartoon representation with the primary monomer shown in green and the dimerization region from the second monomer shown in blue. (E) The Atg11-NTD hybrid structural model from D is shown as a cartoon representation with each domain labeled. The primary monomer is shown in green and the second monomer, which only contains the dimerization region, is shown in blue.

The Atg11-NTD contains a ubiquitin-like domain (ULD) between amino acids 22 and 95. This ULD packs against a scaffold like domain (SLD) spanning amino acids 170 and 455 that connects to a dimerization region between 456 and 596 (**Figure 3E**). The dimerization region of the Atg11-NTD is mediated by a long alpha helix spanning amino acids 456 to 510 and three additional shorter helices all of which have visible density in the reconstruction demonstrating that these regions are ordered in the structure and likely important for stabilizing the dimer interface. The ULD and SLD of the Atg11-NTD superimpose well on the ULD and SLD of human FIP200 (**Figure S4**) [17,20]. However, the dimerization regions of Atg11 and FIP200 are different. Atg11 has a multi helical bundle from each monomer to stabilize dimerization while the dimerization of FIP200 is supported by a single long helix from each monomer. This results in the overall shape of the Atg11-NTD dimer being different from that of FIP200, with the former having a Z shape while the latter has a U shape.

### Atg9(1-255) binding to Atg11-NTD requires a positively charged binding surface

Having demonstrated that the AlphaFold 3 Atg11-NTD prediction is consistent with our low resolution reconstruction, we used AlphaFold 3 to predict where Atg9(1-255) may bind on the surface of the Atg11-NTD. We used a dimer of the Atg11-NTD and one copy of Atg9(1-255) as an input for structure prediction. The top five Atg11-NTD predictions aligned well with very high and confident pLDDT scores for the regions that correspond to the experimental density of the Atg11-NTD cryo-EM structure (**Figure S5A and B**). The Atg11-NTD AlphaFold model has a hydrophobic cavity formed by the ULD sitting against the SLD (**Figure 4A**). The top five models for Atg9(1-255) bound to the Atg11-NTD docked some amino acids from Atg9 into this cavity with the backbone of these Atg9 amino acids superimposing well across the models. This binding region was the only region of structural overlap between all five of the top Atg9(1-255) models (**Figure S5A-C**). The majority of Atg9(1-255) did not overlay at all and had very low pLDDT scores, which indicates poor confidence in the conformation and is likely due to the disordered nature of Atg9(1-255) (**Figure S5B**) [31]. In comparison, the regions that docked into the hydrophobic cavity on Atg11 mostly had low pLDDT scores and a few had confident pLDDT scores (**Figure S5C**). This indicates slightly higher confidence in this region of Atg9 compared to the rest of the structure. Three of the top five predictions, including the top model, docked the first PLF motif (163-165) into this cavity, suggesting that this region may be a possible binding site for Atg9. Despite the primarily low confidence scores for this region, we decided to test if Atg9 may be binding to this region in Atg11 by disrupting the hydrophobic binding pocket by generating an Atg11-NTD (R96D, F99A, Y218A, Y222A, M423A, L426A) mutant construct and purifying it. However, this mutation resulted in recombinant protein that was insoluble, preventing us from testing the importance of this hydrophobic binding pocket in Atg9 binding.

**Figure 4.**
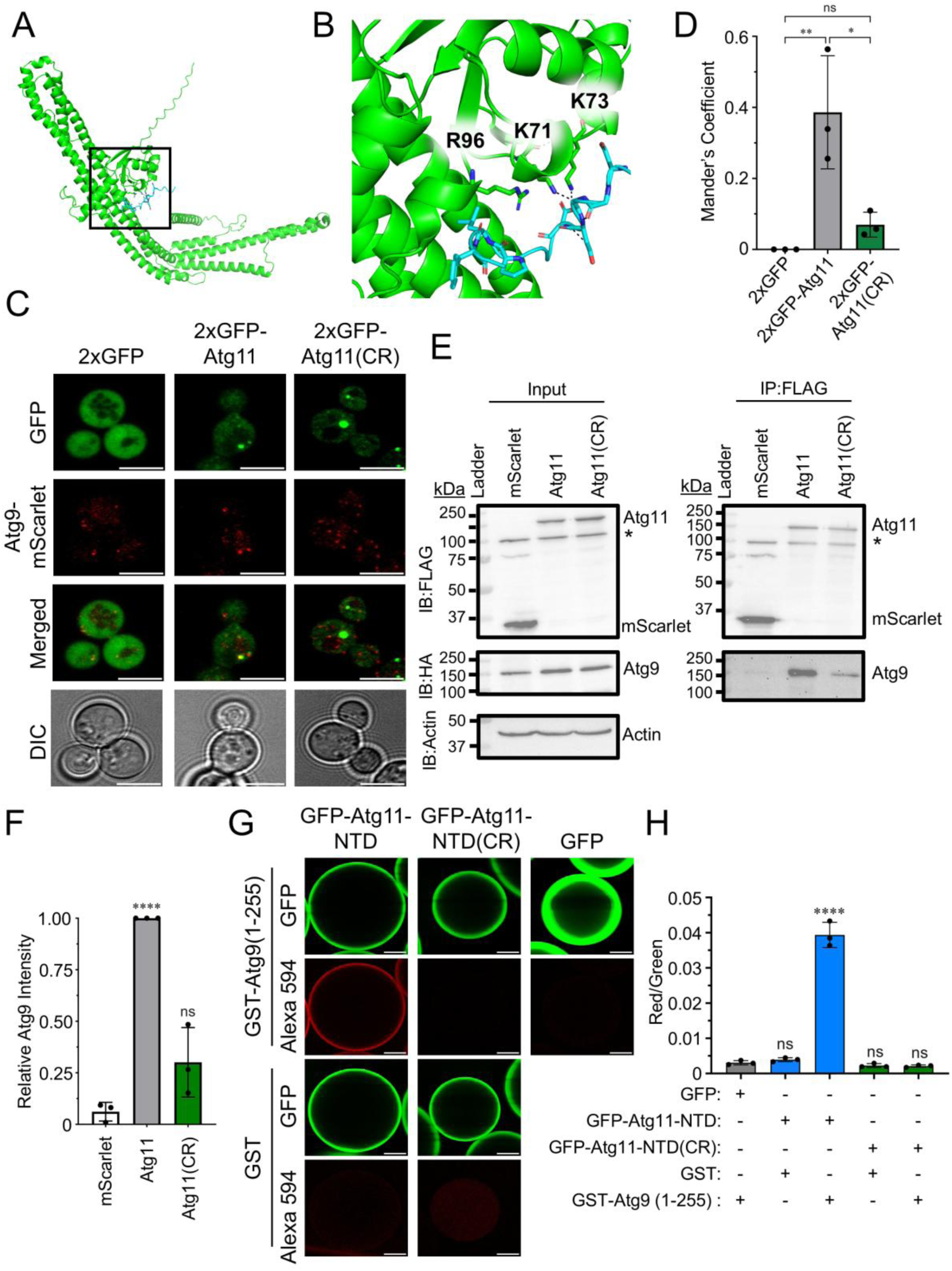
A positively charged binding pocket in the Atg11-NTD is required for Atg9 binding. (A) AlphaFold 3 model of one monomer of the Atg11-NTD dimer shown in green with one copy of Atg9(1-255) shown in blue. Only the amino acids from Atg9 that directly contact Atg11 are shown. The region where Atg9 is predicted to bind to Atg11 is boxed. (B) A close-up view of the boxed area from A. Predicted polar interactions are shown as dashed lines. The three positively charged amino acids that are mutated in the Atg11 CR mutant are shown as stick representations and labeled. (C) Representative microscopy images of *atg11*Δ *atg9*Δ *S. cerevisiae* transformed with the indicated 2xGFP-Atg11 construct and Atg9-mScarlet grown in SMD. Scale bars: 5 μm. (D) Mander’s Coefficient quantification of the images from C. Data represents means ± SD from three independent repeats (n=3). * indicates p ≤ 0.05, ** indicates p ≤ 0.01, ns indicates non-significant calculated using a One-way ANOVA with Tukey’s multiple comparisons test. (E) Representative Western blots of co-immunoprecipitation assays with *atg11*Δ *atg9*Δ *atg17*Δ *S. cerevisiae* transformed with 3xFLAG-mScarlet or the indicated 3xFLAG-Atg11 construct with Atg9-HA grown in SMD. Each blot is labeled with the antibody that was used. Actin was used as a loading control. * indicates non-specific bands. (F) Quantification of the relative Atg9 band intensity from the Western blots in E. Data represents means ± SD from three independent repeats (n= 3). **** indicates p < 0.0001, ns indicates non-significant calculated using Welch’s unpaired t-test. (G) Representative fluorescence microscopy images of GFP-trap beads with the indicated GFP-Atg11-NTD constructs or GFP immobilized then loaded with Alexa594 labeled GST-Atg9(1-255) or Alexa594 labeled GST. Fluorescence intestines were normalized to GFP-Atg11-NTD with GST-Atg9(1-255) image. Scale bars: 25 µm. (H) Quantification of images from G in which fluorescence intensity around the bead was quantified for both green and red channels of 5 beads per images with 3 images per repeat. The ratio from these measurements was then plotted. Data represents means ± SD from three independent repeats (n=3). **** indicates p ≤0.0001, ns indicates non-significant calculated using One-way ANOVA with Tukey’s multiple comparisons test

We also observed that there is a positively charged binding pocket on the surface of Atg11 that is directly adjacent to the hydrophobic pocket predicted to bind to the first PLF motif on Atg9. This positively charged pocket on Atg11 was predicted to coordinate interactions with the negatively charged amino acids preceding the first PLF motif in Atg9 (**Figure 4B and S5D**). Several of the positively charged amino acids in Atg11 that form this pocket, including Lys71, Lys73 and Arg96 are highly conserved in Atg11 across different yeast species suggesting that these amino acids may be important for the function of Atg11 (**Figure S5E**). We hypothesized that if Atg9 is binding to this positively charged pocket on Atg11 then charge reversal mutations in Atg11 may disrupt the interaction between Atg9 and Atg11. Therefore, we generated a triple charge reversal (CR) mutant of Atg11 (K71E, K73E, R96E), referred to as Atg11(CR) for the rest of the manuscript. GFP-Atg11-NTD(CR) could be recombinantly expressed and purified from *E. coli* in a nearly identical fashion to the GFP-Atg11-NTD protein indicating that these mutations do not disrupt the overall fold or stability of the protein. In addition, to further confirm that the CR mutation does not disrupt the Atg11-NTD fold, we performed circular dichroism spectroscopy which revealed that GFP-Atg11-NTD has a similar secondary structure content to GFP-Atg11-NTD (**Figure S6**).

To determine whether Atg11(CR) can interact with Atg9 we first monitored the colocalization of 2xGFP-Atg11 and 2xGFP-Atg11(CR) with Atg9-mScarlet in *atg11Δ atg9Δ S. cerevisiae* (**Figure 4C**). 2xGFP-Atg11(CR) showed reduced colocalization with Atg9-mScarlet with a Mander’s coefficient of 0.07 ± 0.04, contrasting 2xGFP-Atg11 and Atg9-mScarlet with a Mander’s coefficient of 0.39 ± 0.16 (**Figure 4D**). This difference suggests that Atg11 amino acids K71, K73 and R96 are critical for Atg9 recruitment in cells. To verify these findings, we performed co-immunoprecipitation experiments with 3xFLAG-Atg11 or 3xFLAG-Atg11(CR) and Atg9-HA in *atg11Δ atg9Δ atg17Δ S. cerevisiae*. Atg9 co-immunoprecipitated with Atg11 as expected (**Figure 4E**). In contrast, Atg9 co-immunoprecipitation with Atg11(CR) was reduced by approximately 64%, further supporting the important role for these positively charged amino acids in Atg11 for Atg9 binding (**Figure 4F**). Importantly, 3xFLAG-Atg11 and 3xFLAG-Atg11(CR) both expressed at similar levels further supporting that these mutations do not disrupt the stability of Atg11 (**Figure 4E**).

To further explore the impact of the positively charged amino acids in Atg11 on the direct binding interaction between Atg11 and Atg9, we performed the GFP-trap bead assay with GFP-Atg11-NTD and GFP-Atg11-NTD(CR) (**Figure 4G**). The presence of the three CR mutations resulted in a complete loss of binding of Alexa 594 labeled GST-Atg9(2-255) further supporting the important role of these amino acids in Atg9(2-255) binding (**Figure 4H**). Taken together, our cellular and biochemical assays support a role for Atg11 amino acids K71, K73, R96 in binding to Atg9.

### Clustering of Atg11-NTD is required for Atg9-Atg11-NTD colocalization

We have demonstrated that Atg9(2-255) binds to the Atg11-NTD in vitro via our bead assay and BLI **(Figure 2)**. However, in cells the Atg11-NTD does not colocalize well or co-immunoprecipitate with Atg9 **(Figure 1)**. We hypothesized that this may be because the Atg11-NTD does not form as many puncta in cells as Atg11 or the Atg11-CTR which would reduce its interaction with Atg9 due to a loss in avidity. To test this idea, we artificially clustered the Atg11-NTD using the µNS viral capsid protein that self assembles into distinct particles in the yeast cytoplasm [36,37]. We performed a colocalization analysis with 2xGFP-Atg11, 2xGFP-Atg11-NTD, 2xGFP-Atg11-NTD-µNS and Atg9-mScarlet in *atg11Δ atg9Δ S. cerevisiae* cells (**Figure 5A**). Colocalization of Atg9-mScaret and 2xGFP-Atg11-NTD-µNS resulted in near wildtype colocalization with Mander’s coefficients of 0.40 ± 0.07 for 2xGFP-Atg11-NTD-µNS and 0.51 ± 0.16 for 2xGFP-Atg11. In contrast, 2xGFP-Atg11-NTD had essentially no colocalization with Atg9-mScarlet with a Mander’s coefficient of 0.005 ± 0.004, consistent with our previous observations (**Figure 5B and 1E**). We next investigated if the recruitment of Atg9 to Atg11-NTD-µNS is dependent on the positively charged binding pocket on the Atg11-NTD. For this we performed a colocalization analysis of 2xGFP-Atg11-NTD-µNS or 2xGFP-Atg11-NTD(CR)-µNS with Atg9-mScarlet (**Figure 5C**). 2xGFP-Atg11-NTD(CR)-µNS was still able to form puncta, but had reduced colocalization with Atg9-mScarlet when compared to 2xGFP-Atg11-NTD-µNS, with Mander’s coefficients of 0.032 ± 0.006 and 0.23 ± 0.015, respectively (**Figure 5D**). These data strongly support the need for the positively charged binding pocket on Atg11-NTD to facilitate an Atg11-NTD-Atg9 interaction which also requires the clustering of the Atg11-NTD by the CTR for their complete interaction in cells.

**Figure 5.**
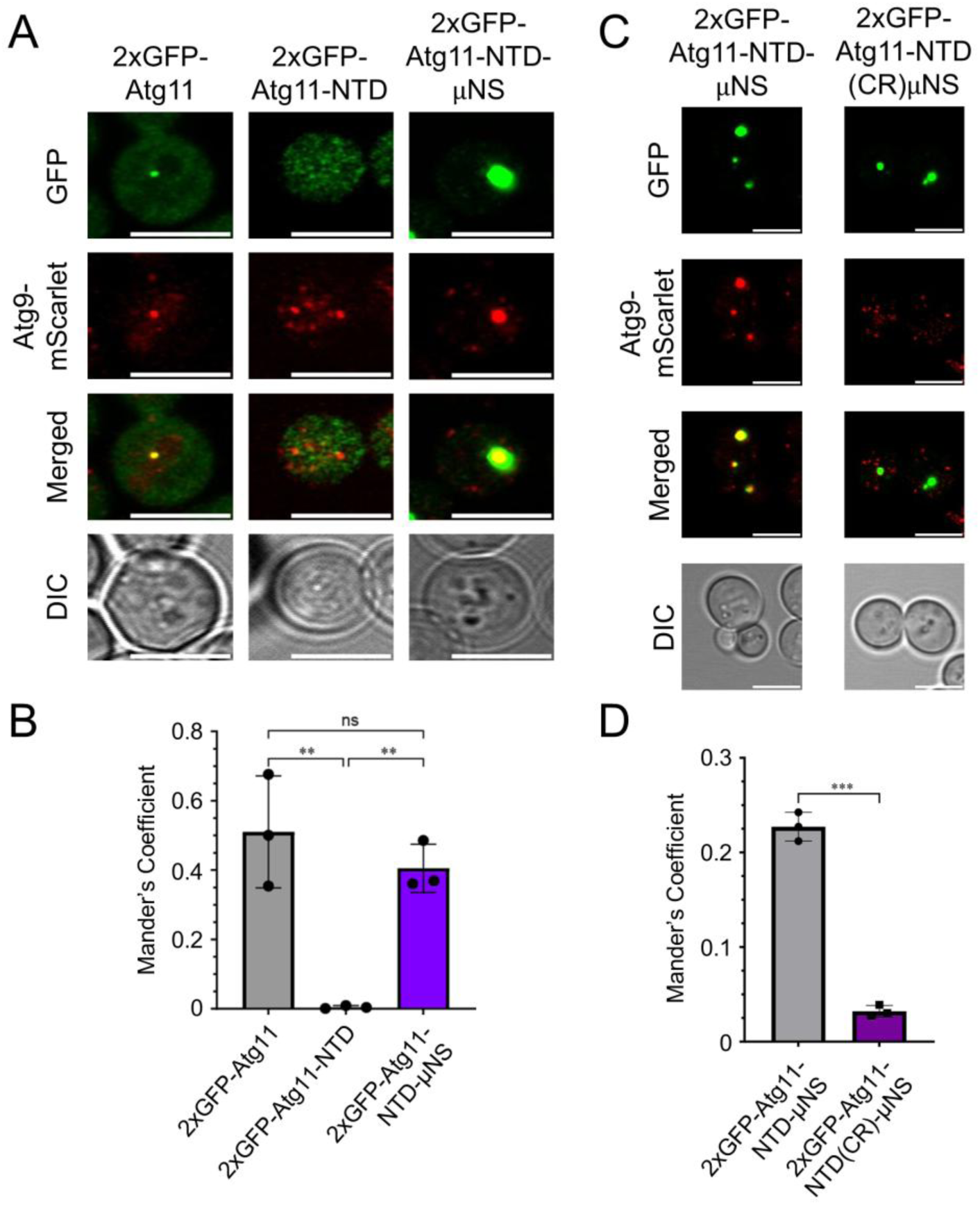
Clustering of the Atg11-NTD facilitates its colocalization with Atg9. (A) Representative microscopy images of *atg11*Δ *atg9*Δ *S. cerevisiae* transformed with the indicated 2xGFP-Atg11 construct and Atg9-mScarlet grown in SMD. Scale bars: 5 μm. (B) Mander’s Coefficient quantification of the images from A to display the co-occurrence of Atg9-mScarlet with the indicated 2xGFP-Atg11 constructs. Data represents means ± SD from three independent repeats (*n*= 3). ** indicates p ≤ 0.01, ns indicates non-significant calculated using a One-way ANOVA with Tukey’s multiple comparisons test. (C) Representative microscopy images of *atg11*Δ *atg9*Δ *S. cerevisiae* transformed with the indicated 2xGFP-Atg11 constructs and Atg9-mScarlet grown in SMD. Scale bars: 5 μm. (D) Mander’s Coefficient quantification of the images from C. Data represents means ± SD from three independent repeats (*n*= 3). *** indicates p ≤ 0.001, calculated by Welch’s unpaired t-test.

### The Atg11 CR mutation leads to a reduction in mitophagy

To test if the Atg11 CR mutation influences selective autophagy in cells, we monitored the progression of mitochondrial autophagy (mitophagy) using three complimentary yeast mitophagy assays. Atg32 is the transmembrane selective autophagy receptor for mitophagy in yeast [38,39]. Under basal conditions, Atg32 has a diffuse localization in the outer mitochondrial membrane (OMM). Once mitophagy is induced, Atg11 binds to Atg32, leading to a rapid clustering of Atg32 in the OMM, occurring as one of the first steps in mitophagy [39,40]. To determine if the Atg11(CR) mutation impacts Atg32 clustering during mitophagy initiation, we transformed *atg11Δ S. cerevisiae* with either 3xMyc-Atg11 or 3xMyc-Atg11(CR) and GFP-Atg32. Cells were grown in nutrient rich media, then shifted to nitrogen starved media for 15-30 minutes to induce mitophagy (**Figure 6A**). GFP-Atg32 puncta were reduced in the presence of 3xMyc-Atg11(CR) compared to 3xMyc-Atg11, with 17.7 ± 7.32% and 45.5 ± 11.4% of cells having Atg32-GFP puncta, respectively (**Figure 6B**). This suggests that the Atg11 CR mutation results in a reduction in one of earliest events during mitophagy initiation and may suggest a correlation between Atg9 recruitment and Atg32 clustering.

**Figure 6.**
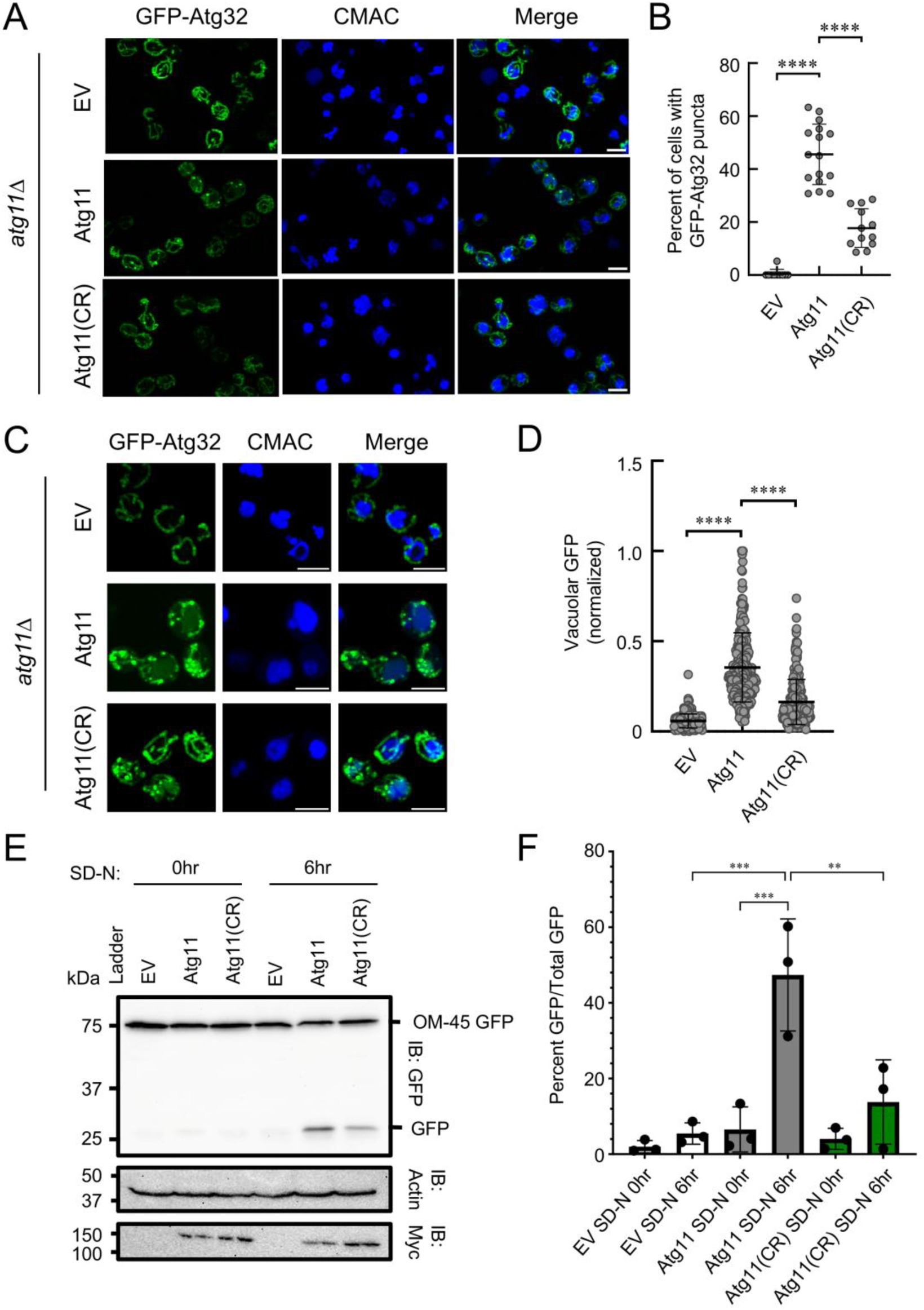
A positively charged binding pocket on Atg11 is required for mitophagy. (A) Representative microscopy images *atg11Δ S. cerevisiae* cells transformed with GFP-Atg32 and empty vector (EV), Atg11 or Atg11(CR) were subjected to nitrogen starvation media (SD-N). Cells were imaged between 15 and 30 minutes following starvation. Vacuoles were stained with CMAC. Scale bar = 5 μm. (B) Quantification of the images from A showing the percentage cells with at least one GFP-Atg32 punctum. The graph shows pooled data from three independent repeats (n=3). Each point represents quantification of all yeast cells from a single field. Between 3 and 6 fields were analyzed per condition per repeat. **** indicates p≤0.0001 analyzed using an ordinary one-way ANOVA with Šídák’s multiple comparisons test. (C) Representative microscopy images of *Δatg11 S. cerevisiae* cells expressing GFP-Atg32 transformed with empty vector (EV), Atg11 or Atg11(CR) were subjected to nitrogen starvation for 90 minutes and imaged. Vacuoles were stained with CMAC. Scale bar = 5 μm. (D) Quantitation of vacuolar GFP intensities from C. Graph shows pooled data from three independent repeats. Each data point represents GFP intensity from a single vacuole. At least 274 vacuoles were analyzed per condition. **** indicates p≤0.0001 analyzed using Ordinary one-way ANOVA with Šídák’s multiple comparisons test. (E) Representative western blots of an OM45-GFP assay in which *atg11*Δ *S. cerevisiae* cells expressing OM45-GFP were transformed with empty vector (EV), 3xMyc-Atg11, or 3xMyc-Atg11(CR). Cells were then grown in lactate media for 14-16 hours and then shifted to nitrogen starvation media (SD-N) to induce mitophagy. Actin was used as a loading control. Each blot is labeled with IB indicating what antibody was used. (F) Quantification of the Western blots from E in which GFP intensity divided by OM-45 GFP intensity is plotted as a percentage. Data represents means ± SD from three independent repeats (*n*= 3). *** indicates p ≤ 0.001, ** indicates p ≤ 0.01, n.s. indicates non-significant analyzed using an Ordinary one-way ANOVA with Šídák’s multiple comparisons test.

As mitophagy proceeds, mitochondria containing GFP-Atg32 puncta are fissioned from the mitochondrial network, captured in autophagosomes and delivered to the vacuole for degradation [41,42]. In the vacuole, Atg32 is quickly degraded while GFP persists. As a result, the amount of GFP in the vacuole is proportional to the amount of Atg32 targeted to the vacuole during mitophagy. Therefore, we next investigated if mitophagy completion is impacted by the Atg11(CR) mutant by inducing mitophagy and measuring the amount of GFP in the vacuole after 30 minutes (**Figure 6C**). The amount of GFP found in the vacuoles was reduced by 46% in the presence of 3xMyc-Atg11(CR) when compared to the average amount of GFP found in vacuoles when 3xMyc-Atg11 is present (**Figure 6D**). This suggests that mitophagy is reduced in the presence of Atg11(CR) but not completely blocked.

To verify our results demonstrating that Atg11(CR) leads to reduced mitophagy we used the Om45-GFP reporter assay [43]; [44]. In this assay an outer mitochondrial membrane protein, Om45, is fused to GFP on its C-terminus. Upon completion of mitophagy, Om45 is degraded in the vacuole, but GFP persists. This produces an amount of free-GFP that is proportional to the amount of mitochondria captured in the vacuole. Since Atg32 serves as the receptor and OM45 as a mitochondrial cargo, both the GFP-Atg32 and OM45-GFP assays are complimentary. We transformed *atg11*Δ cells containing OM45-GFP with empty vector, 3xMyc-Atg11 or 3xMyc-Atg11(CR). Western blots showed that in the presence of 3xMyc-Atg11(CR), the amount of free-GFP was reduced by 71% indicating that mitophagy is hindered in the presence of the CR mutation but not completely blocked (**Figure 6E and 6F**). This is in excellent agreement with our GFP-Atg32 results and further confirms that the positively charged binding pocket on Atg11 is critical for selective autophagy. Importantly, 3xMyc-Atg11 and 3xMyc-Atg11(CR) expressed at similar levels in yeast indicating that the reduction in mitophagy is not due to reduced protein levels as a result of the CR mutation (**Figure 6E**). Taken together, these results demonstrate that the CR mutation leads to a reduction in mitophagy that can be observed at even the earliest stages of mitophagy initiation.

## DISCUSSION

In this paper we demonstrate that the N-terminal disordered region of Atg9 interacts directly with the Atg11-NTD via the previously identified PLF motifs in Atg9 [31]. We demonstrate that clustering of the Atg11-NTD by the CTR is required in cells to bind Atg9, as the Atg11-NTD expressed by itself is unable to colocalize or immunoprecipitate with Atg9. However, Atg11-NTD binding to Atg9 could be recovered in cells when the Atg11-NTD was clustered artificially by the µNS. We analyzed the structure of the Atg11-NTD using cryo-EM which revealed a Z shaped dimeric structure which is consistent with the AlphaFold model of dimeric Atg11-NTD. The Atg11-NTD contains a ULD packed against an SLD which is similar to FIP200 but Atg11 contains a dimerization region that differs from FIP200, giving the two proteins a distinctly different overall shape [17,20]. The ULD and SLD of the Atg11-NTD form an extended binding cavity that was predicted to interact with Atg9 using AlphaFold 3. Mutation of the positively charged surface of this binding pocket led to a loss in Atg9 binding in vitro and in vivo and resulted in a reduction in mitophagy, altogether demonstrating the importance of this positively charged binding pocket.

Atg11 was originally demonstrated to interact with Atg9 via the Atg11-NTD and not the CTR using yeast two-hybrid assays [30]. However, our work suggests that the interaction between the Atg11-NTD and Atg9 is sufficiently weak in cells without the CTR that it is difficult to detect by both colocalization and immunoprecipitation assays. We also demonstrate that the CTR can be replaced with another clustering domain (µNS) to fully recover the interaction between Atg11 and Atg9. This suggests that the CTR itself is not required for Atg9 binding but instead drives clustering of Atg11 which is essential to stabilize the interaction between Atg11 and Atg9 in cells. This observation that the CTR is required for Atg11 to bind to Atg9 is also supported by earlier work which showed a significant reduction in co-immunoprecipitation between Atg11 and Atg9 when the CTR was removed [45]. Our in vitro binding data further supports that the interaction between Atg11 and Atg9 is weak, with a 4.63 ± 0.757 µM K_d_, and that the interaction is stabilized in cells through the enhanced avidity of many copies of Atg11 being brought together in close proximity to multiple copies of Atg9. The importance of avidity for protein interactions has now become a recurring theme in autophagy, whereby weak interactions between autophagy proteins do not fully support stable 1 to 1 interactions in cells. Instead, oligomerization, clustering or phase separation are critical for these protein interactions to be stabilized in cells [36,46–48].

Previous work demonstrated that Atg17 and Atg11 compete for binding to Atg9 and that binding of Atg32 activates Atg11 to bind to Atg9 [34]. This suggests that SAR binding to Atg11 is a requirement for Atg9 binding. However, our results demonstrate that SAR binding to Atg11 is not needed for Atg11 to interact with Atg9 since replacing the entire CTR with µNS is sufficient to trigger colocalization between Atg11 and Atg9 in cells with similar Mander’s coefficients between full-length Atg11 and Atg9 and Atg11-NTD-µNS. One effect of SAR binding to Atg11 is that it leads to the clustering of Atg11. For example, when Atg11 binds to Atg32 in the outer mitochondrial membrane Atg32 and Atg11 are rapidly clustered into puncta [19,40,49]. Similarly, Atg11 binding to Atg19, another SAR, leads to the clustering of Atg11 at the large oligomeric APE1 cargo [14,50]. In both cases, SAR binding to Atg11 results in clustering of Atg11 which would effectively enhance the interaction strength between Atg11 and Atg9 leading to a stable interaction between these binding partners. Therefore, one possible mechanism by which SARs drive the interaction between Atg11 and Atg9 may be partially through the clustering of Atg11.

We also reveal a novel potential protein binding surface on Atg11 comprised of K71, K73, R96. We demonstrate that when this positively charged binding pocket is mutated to negatively charged amino acids, Atg9 binding is lost both in vitro and in cells. This loss of binding strongly supports a role for these amino acids in the interaction of Atg11 and Atg9. Intriguingly, the AlphaFold model of the Atg11-NTD shows that this binding pocket in Atg11 is formed by the ULD packing against the SLD. This specific region of Atg11 has not been directly attributed to protein binding previously. Previous attempts to narrow down the Atg9 binding site on Atg11 identified residues 537-576 as being required [50]. Our low resolution cryo-EM volume combined with AlphaFold modeling reveals that 537-576 resides in the middle of the dimerization domain of Atg11-NTD. Therefore, it is possible that disrupting this region of the protein may destabilize the overall fold of the Atg11-NTD thus blocking Atg9 binding. In addition, the Backues group demonstrated that the Atg11 Y565E mutation resulted in reduced Atg9 interaction via yeast-2-hybrid and co-immunoprecipitation assays [51]. They also found that this residue was required for Atg11 self-interaction which made them suspect that this may be impacting the structure and stability of Atg11 [51]. Their data are consistent with our prediction that the dimerization region of the Atg11-NTD is required for the overall stability of the Atg11-NTD. Lastly, Atg11 I569 was found to also facilitate Atg9 binding via yeast-2-hybrid and co-immunoprecipitation during glucose-starved autophagy [52]. I569 lies directly in the dimerization of the Atg11-NTD. Consequently, its mutation may disrupt dimerization, and thus the overall stability of the Atg11-NTD. In contrast, we demonstrated that the mutation of K71, K73 and R96 does not directly disrupt the overall fold or stability of the Atg11-NTD and instead lead to a loss in Atg9 binding.

We obtained a 6.39 Å cryo-EM reconstruction of dimeric Atg11-NTD which we used with local refinement and AlphaFold modeling to model one copy of the Atg11-NTD along with the dimerization region from the second copy of Atg11. Our hybrid model reveals that human FIP200 contains a similar ULD and SLD fold but that the dimerization regions of these proteins are distinct [20]. The ULD and SLD in FIP200 form a similar positively charged binding pocket surrounding K29, but the pocket is less charged than the pocket in Atg11. The dimerization domain of the FIP200-NTD consists of a single long alpha helix from each monomer which causes the protein to form a U shape. In contrast, the Atg11-NTD dimerization domain consists of a longer primary alpha helix, supported by three additional shorter helices. This enables Atg11 to form an elongated Z shape giving a different overall shape from FIP200. We previously reported that Atg11 contains a predicted amphipathic helix between amino acids 612 and 646, which we have shown is important for Atg11 membrane binding [32]. Our cryo-EM reconstruction does not contain any density for this region of the structure supporting the idea that this region is dynamic and not part of the overall Atg11-NTD fold and therefore available for membrane binding.

Several cryo-EM structures of FIP200 have been determined in complex with its binding partners, including ULK1, Atg13, and the PI3K complex 3 [17,20]. These structures all have one thing in common, the positively charged binding pocket formed by the ULD and SLD in FIP200 is empty. Therefore, if FIP200 binds to Atg9A, the mammalian counterpart of Atg9, via a similar pocket as Atg11 then it is possible that Atg9A may be able to bind into this binding pocket when FIP200 is bound to its other protein interaction partners including ULK1 and Atg13. At this time, it is unclear if Atg11 binds to Atg1 and Atg13 with a similar structure to FIP200, but if this is the case, the positively charged binding pocket would potentially support Atg9 binding simultaneously with Atg1 and Atg13.

## MATERIALS AND METHODS

### Plasmid Generation

Plasmids used in this study are listed in Table S2 and were generated using New England Biolabs Q5 site directed mutagenesis (SDM) kit, Gibson Assembly (GA) or were purchased. All plasmids were verified by Sanger or plasmid sequencing. Plasmids pSN040, pSN054, pSK073 were generated by SDM or GA into pET His6 TEV LIC cloning vector (1B) which was a gift from Scott Gradia (Addgene plasmid # 29653). Plasmids pSN074 and pSN021 were generated by SDM or GA into pET His6 GFP TEV LIC cloning vector (1GFP) which was a gift from Scott Gradia (Addgene plasmid # 29663). Plasmid pSN060 was generated by TWIST Bioscience where the Atg11-NTD was codon optimized for *E. coli* expression and subcloned into a pET21 vector. Plasmid pZB036 was generated by SDM using restriction enzyme sites NcoI and SalI to insert Atg9(1-255) into pET-22A vector. Plasmid pZB051 was generated using GA into the pET-22A vector. Plasmids pSN007, pSN008, pSN010, pSN019, pHMP001, pZG001 and pSN069 were generated by SDM or GA in respective yCPLac111 or yCPlac33 vector [53]. Plasmid pSN080 was generated by Genscript. The GFP-Atg32 plasmid was a generous gift from Dr. Benedikt Westermann [54]. Plasmid pDA078 was generated by TWIST biosciences by inserting 3xFLAG-mScarlet into the pCu415 vector [55]. Plasmid pZB078 was generated using GA into yCPlac111.

### Protein Expression and Purification

#### GFP-Atg11-NTD & GFP-Atg11-CTR

6xHis-GFP-TEV-Atg11-NTD (1-646) and 6xHis-GFP-TEV-Atg11-CTR (698-1178) were both recombinantly expressed and purified from *E. coli* using the same procedure. The relevant plasmid was transformed into BL21 (DE3) Rosetta2 cells and grown at 37 ^°^C to and OD_600_ of ∼0.6 then induced with 0.1 mM IPTG and grown at 18 ^°^C overnight. Cells were then harvested by centrifugation at 4000 x g at 4 ^°^C for 25 minutes. Harvested cells were resuspended in Atg11 lysis buffer (50 mM Tris pH 8.0, 500 mM NaCl, 5 mM MgCl2, 1% [v/v] Triton X-100, 1 mM PMSF and a Roche cOmplete EDTA free tablet) and lysed by passing through a microfluidizer at 18,000 psi three times. Lysate was then cleared by centrifugation at 40,000 x g for 45 minutes at 4 ^°^C. Cleared supernatant was then added to TALON resin that was pre-equilibrated with Atg11 lysis buffer. Talon resin was then washed three times with 50 mM Tris pH 8.0, 300 mM NaCl and then once with 50 mM Tris pH 8.0, 300 mM NaCl, 2.5 mM imidazole. Protein was then eluted with 50 mM Tris pH 8.0, 300 mM NaCl, 200 mM imidazole. Eluates containing protein were verified by SDS-PAGE and then concentrated and subjected to size exclusion chromatography (SEC) using a HiLoad 16/60 Superdex 200 PG equilibrated with 20 mM Tris pH 8.0, 300 mM NaCl, 0.2 mM TCEP. Samples with GFP-Atg11-NTD or GFP-Atg11-CTR were verified by SDS-PAGE, pooled and concentrated to 10-15 µM and frozen using liquid nitrogen and stored at −80 ^°^C until used. Before any protein was used for bead assays, protein was dialyzed into buffer containing 20 mM Tris pH 8.0, 150 mM NaCl, 0.2 mM TCEP.

#### GST-Atg9(1-255)

Purification of GST-Atg9(1-255) was performed as described previously [56]. Briefly, 10xHis-GST-TEV-Atg9(1-255) was transformed into BL21 Rosetta2 (DE3) cells and grown in LB with 2 g/L glucose to suppress initial expression. Cells were grown to an OD_600_ of ∼2 at 37 ^°^C then induced with 0.1 mM IPTG and grown at 37 ^°^C for 2 hours. Cells were then harvested, resuspended in Atg9 lysis buffer (20 mM sodium phosphate pH 7, 500 mM NaCl,1 mM PMSF) and lysed by passing through a microfluidizer at 18,000 psi three times. Lysate was then cleared by centrifugation at 55,000 x g for 20 minutes at 4 ^°^C. Cleared supernatant was initially purified using TALON resin equilibrated with Atg9 lysis buffer. Resin was then washed with 20 mM sodium phosphate pH 7, 500 mM NaCl,1mM PMSF then with 20 mM Tris pH8, 150 mM NaCl, 2.5 mM imidazole. Protein was eluted with 20 mM Tris pH8, 150 mM NaCl, 150 mM imidazole. Eluates were assessed by SDS-PAGE and fractions containing protein were pooled, concentrated and purified further using SEC with a HiLoad 16/60 Superdex 200 PG column equilibrated with 20 mM Tris pH 8.0, 150 mM NaCl, 0.2 mM TCEP. Fractions were evaluated using SDS-PAGE and samples containing 10xHis-GST-TEV-Atg9(1-255) were concentrated to 40 µM and frozen using liquid nitrogen and stored at −80 ^°^C until used.

#### Atg11-NTD for BLI and Cryo-EM

Atg11-NTD-TEV-6xHis was codon optimized for expression in *E. coli* by TWIST Bioscience and subcloned into the pET21 vector. The plasmid was transformed into BL21 (DE3) STAR cells and grown at 37 ^°^C to an OD_600_ of ∼0.6 then induced with 1 mM IPTG and grown at 18 ^°^C overnight. Cells were harvested at 4000 x g at 4 ^°^C for 25 minutes and resuspended in Atg11 BLI lysis buffer containing (50 mM Tris pH 8.0, 500 mM NaCl, 5 mM MgCl2, 1% [v/v] Triton X-100, 1 mM PMSF and one Roche cOmplete EDTA free tablet). Cells were lysed by three passes through a microfluidizer at 18,000 psi. Lysate was then cleared by centrifugation at 40,000 x g for 45 minutes at 4 ^°^C. Supernatant was initially purified using TALON resin which was equilibrated with Atg11 BLI lysis buffer. Resin was then washed 3 times with 50 mM Tris pH 8.0, 300 mM NaCl and then once with 50 mM Tris pH 8.0, 300 mM NaCl, 2.5 mM imidazole. Protein was then eluted with 50 mM Tris pH 8.0, 300 mM NaCl, 200 mM imidazole. Protein was evaluated by SDS-PAGE. Fractions containing protein were pooled and buffer exchanged into 50 mM Tris pH 8.0, 300 mM NaCl using a HiPrep 26/10 desalting column. Protein was mixed with TEV protease at 1:25 overnight at 4 ^°^C. The sample was then repurified using TALON resin to separate the cleaved and uncleaved protein samples. Flowthrough from the second TALON resin was pooled and further purified using SEC with a HiLoad 16/60 Superdex 200 PG equilibrated in buffer consisting of 20 mM Tris pH 7.4, 150 mM NaCl, 0.2 mM TCEP. Fractions were evaluated by SDS-PAGE and fractions containing Atg11-NTD were pooled and verified to be lacking a His tag via western blot then concentrated and frozen using liquid nitrogen and stored at −80 ^°^C until used.

#### 6xHis-Atg9(1-255) and 6xHis-Atg9(1-255ΔPLF) for BLI

6xHis-Atg9(1-255)-TEV-MBP and 6xHis-Atg9(1-255ΔPLF)-TEV-MBP plasmids were expressed and purified using the same procedure as follows. The plasmid was transformed into BL21 (DE3) STAR cells and grown to and OD_600_ of ∼0.6 at 37 ^°^C then induced with 1 mM IPTG and grown at 18 ^°^C overnight. Cells were then harvested by centrifugation at 4000 x g for 25 minutes at 4 ^°^C. Cells were resuspended in Atg9 BLI lysis buffer (20 mM Tris pH 7.4, 300 mM NaCl, 2 mM MgCl2, 1 mM PMSF, one Roche cOmplete EDTA free tablet) and lysed by three passes through a microfluidizer at 18,000 psi. Lysate was then cleared by centrifugation at 40,000 x g for 30 minutes at 4 ^°^C. Cleared supernatant was initially purified using TALON resin equilibrated with Atg9 BLI lysis buffer. Resin was washed three times with 20 mM Tris pH 8.0, 300 mM NaCl and then once with 20 mM Tris pH 8.0, 300 mM NaCl, 2.5 mM imidazole. Protein was eluted with 20 mM Tris pH 8.0, 300 mM NaCl, 200 mM imidazole and fractions were evaluated using SDS-PAGE. Fractions containing protein were pooled and buffer exchanged into 20 mM Tris pH 8.0, 300 mM NaCl using a HiPrep 26/10 desalting column. Protein was then mixed with TEV protease at 1:25 overnight. Since Atg9 (1-255) is intrinsically disordered and vulnerable to proteolysis the next day the sample was heated at 95 ^°^C for 15 minutes and then centrifuged at 20,000 x g for 5 minutes to inactivate and precipitate structured proteins. Supernatant was then concentrated and further purified using SEC with a HiLoad 16/600 Superdex 200 PG column equilibrated in buffer consisting of 20 mM Tris pH 7.4, 150 mM NaCl, 0.2 mM TCEP. Samples were evaluated using SDS-PAGE and fractions containing Atg9 (1-255) sample were pooled, concentrated, frozen using liquid nitrogen and stored at −80 ^°^C until used.

#### mScarlet-Atg9(1-255) and mScarlet-Atg9(1-255)ΔPLF

TwinStrep-mScarlet-Atg9(1-255)-trifold-6xHis and TS-mScarlet-Atg9(1-255ΔPLFs)-trifold-6xHis plasmids were purified using the same procedure as follows. The plasmid was transformed into BL21 (DE3) STAR cells and grown to and OD_600_ of ∼0.6 at 37 ^°^C then induced with 0.5 mM IPTG and grown at 18 ^°^C overnight. Cells were then harvested by centrifugation at 4000 x g for 25 minutes at 4 ^°^C. Cells were resuspended in mScarlet-Atg9 lysis buffer (20 mM Tris pH 8, 300 mM NaCl, 2 mM PMSF, one Roche cOmplete EDTA free tablet) and lysed by three passes through a microfluidizer at 18,000 psi. Lysate was then cleared by centrifugation at 40,000 x g for 45 minutes at 4 ^°^C. The supernatant was then added to a column containing TALON resin equilibrated in mScarlet-Atg9 lysis buffer. Resin was washed three times with 20 mM Tris pH 8.0, 300 mM NaCl and once with 20 mM Tris pH 8.0, 300 mM NaCl, 2.5 mM imidazole. Protein was then eluted with 20 mM Tris pH 8.0, 300 mM NaCl, 200 mM imidazole. Samples were evaluated using SDS-PAGE and fractions containing protein were pooled and further purified using Strep-Tactin XT 4 flow high-capacity resin equilibrated in 20 mM Tris pH 8.0, 300 mM NaCl. Resin was then washed five times with 20 mM Tris pH 8.0, 300 mM NaCl. Protein was then eluted with 1x BXT (100mM Tris-Cl, 150 mM NaCl, 1 mM EDTA, 50 mM biotin, pH 8). Fractions were evaluated by SDS-PAGE and fractions containing protein were pooled, concentrated and further purified using SEC with a HiLoad 16/600 Superdex 200 PG column equilibrated in 20 mM Tris pH 8.0, 150 mM NaCl, 0.2 mM TCEP. Fractions were evaluated using SDS-PAGE and fractions containing protein were pooled, concentrated, frozen using liquid nitrogen and stored at −80 ^°^C until used.

### Yeast cell lines

All yeast cell lines used in this study are described in Table S3.

### Colocalization Studies and Analysis

*atg11*Δ *atg9*Δ *S. cerevisiae* cells were grown in YPD (1% yeast extract, 2% tryptone, and 2% glucose) at 30 ^°^C to an OD_600_ of 0.8-1 and transformed with the indicated plasmids via the LiAc method and plated on SMD (0.67% yeast nitro-gen base, 2% glucose and supplemented with the appropriate amino acid drop out mixture) plates [57]. Plates were incubated for 3-4 days at 30 ^°^C. After 3-4 days, overnight cultures were started in 3 mL of liquid SMD media. The following day cells were diluted to an OD_600_ of 0.3 in SMD and allowed to grow for approximately 2 hrs. Cells were added to MaTek dishes which had a thin layer of 0.1 mg/ml Concanavalin A solution to allow yeast cells to adhere to the dish. Cells were then imaged on a Zeiss Airyscan LSM 880 microscope using a 63x oil immersion lens. GFP was excited with a 488nm argon laser, mScarlet was excited with a DPSS 561nm laser. A total of 15 images through a Z stack were taken at 0.2 µm apart for a minimum of 3 fields per sample per repeat. Images were then Airyscan processed. For analysis a minimum of 3 fields per repeat were selected and max projected. FIJI plugin JaCOP v2.1.4 was used with default parameters selected, aside from manually thresholding for puncta, was used to determine the Mander’s coefficient [58].

### Puncta per Cell Analysis

*atg11*Δ *S. cerevisiae* cells were grown in YPD to an OD_600_ of 0.8-1 and transformed with the indicated plasmids via the LiAc method and plated on SMD plates containing the appropriate selection [57]. Plates were incubated at 30 ^°^C for 3-4 days. Following 3-4 days overnight cultures were started in liquid SMD media. The following day cells were diluted to an OD_600_ of 0.3 in SMD and allowed to grow for approximately 2 hr. Cells were added to MaTek dishes which had a thin layer of 0.1 mg/ml Concanavalin A solution to allow yeast cells to adhere to the dish. Cells were then imaged on Zeiss Airyscan LSM 880 microscope using a 63x oil immersion lens. GFP was excited with a 488nm argon laser. A total of 15 Z stack images separated by 0.2 µm were taken for a minimum of 3 fields per sample per repeat. Images were then Airyscan processed. For quantification of puncta images were opened in FIJI (ImageJ2) v 2.16.0 and max projected [59]. Cells with visible puncta were counted with a marker as having a puncta and all visible green fluorescence cells were counted as a cell. Puncta counted were determined by drawing a line through the puncta to determine the fluorescence intensity was at least 2 times that of cell fluorescence.

### Co-immunoprecipitation

*atg11*Δ *atg9*Δ *atg17*Δ *S. cerevisiae* cells were grown in YPD to an OD_600_ of 0.8-1 and transformed with the indicated plasmids via the LiAc method and plated on SMD plates containing the appropriate selection [57]. Plates were incubated for 3-4 days at 30 ^°^C. Cells were then grown in 100 mL of SMD to and OD_600_ of 0.8-1.5 and an equal amount of cells for all samples were harvested. Cells were then washed with 1xPBS and resuspended in ten times the OD_600_ equivalent of CO-IP buffer (25 mM Tris-HCl pH 7.4, 150 mM NaCl, 0.2% NP-40, 3 mM PMSF, 5 mM EDTA, and one cOmplete mini protease inhibitor cocktail tablet EDTA-free). This resuspension was placed in tubes and bead-beated at 4 ^°^C twice for 45 seconds with a 5-minute break in between. Samples were centrifuged for 5 minutes at 5000 x g a 4 ^°^C. A sample of the supernatant was saved as “input” and the rest was added to magnetic FLAG beads equilibrated with CO-IP buffer and incubated with end-over-end rocking for 1 hour at 4 ^°^C. Beads were then washed three times with CO-IP buffer and 1x sample buffer was added to the resin at 7.5x of sample buffer to beads volume and heated at 95 ^°^C for 5 minutes and saved as “IP” sample. Samples were then analyzed by Western blot. For HA blots the primary antibody was rabbit-anti-HA antibody from (Cell Signaling Technology: C29F4) at 1:1000 and the secondary antibody was goat anti-Rabbit HRP (Bio-Rad: 1706515). Both were used in 5% milk in 0.1% TBST (Tris-buffered saline, 0.1% [v/v] Tween-20). For FLAG blots the primary was anti-FLAG-HRP from (Millipore Sigma: A8592) at 1:1000 in TBS without a secondary antibody. For actin blots the primary antibody was anti-actin (Thermo Fisher Scientific: MA1-744) at 1:5000 and the secondary antibody was anti-mouse-HRP (Millipore Sigma: A4416) at 1:20000 both in 5% milk in 0.1% TBST.

### Bead Assay and Analysis

GFP-trap beads were generated as follows. 6xHis-TEV-GFP nanobody was transformed into Shuffle T7 Express cells and grown to an OD_600_ of 1 at 37 ^°^C [60]. Cells were induced with 0.1 mM IPTG and grown at 18 ^°^C for 18 hours. Cells were then harvested and lysed in buffer consisting of 20 mM Tris pH 8, 500 mM NaCl. Lysate was clarified and protein was purified using TALON resin. Eluate from the talon resin was further purified using SEC with a Superdex75 16/600 column that was equilibrated in 20 mM phosphate buffered saline pH 7.4, 150 mM NaCl. Sodium bicarbonate was added to a final concentration of 100 mM to size exclusion fractions containing protein. Protein was added to 10 mL of NHS-activated Sepharose 4FF that was equilibrated in 1 mM HCl and incubated overnight at 4 ^°^C. Resin was thoroughly washed with 20 mM Tris pH 7.4. 150 mM NaCl and stored with 0.05% sodium azide at 4 ^°^C until needed.

For the protein-protein binding studies using the GFP-trap beads all protein samples were dialyzed into 20 mM Tris pH7.4, 150 mM NaCl, 0.2 mM TCEP prior to use. GFP-trap beads were equilibrated with GFP-trap wash buffer (20 mM Tris pH 7.4, 150 mM NaCl, 0.2 mM TCEP). 10 µl of beads were loaded with enough GFP tagged protein of interest to saturate a previously determined 1.2mg/ml binding capacity and allowed to incubate for 30 minutes at 4 ^°^C with end-over-end rocking. Beads were then washed three times with GFP-trap wash buffer to remove excess protein. Equimolar amount of non-GFP tagged protein was added to beads and allowed to incubate for 30 minutes at 4 ^°^C with end-over-end rocking. Beads were then washed 3 times with GFP-trap wash buffer to remove excess protein. Beads were then resuspended with 90 µl of GFP-trap wash buffer to create a 10% slurry mix that was imaged on Zeiss Airyscan LSM 880 microscope using a 10x lens. GFP was excited with a 488nm argon laser and mScarlet/Alexa 594 was excited with DPSS 561 laser. For analysis, FIJI (ImageJ2) v 2.16.0 analysis software was used in which the background was subtracted from images in both fluoresce channels, a line was drawn around the bead the intensity in both the 488 and 561 channel was measured. This was repeated for 5 beads per image with 3 images per repeat for a total of 15 beads analyzed per experiment. The ratio was then taken for statistical analysis to be performed in GraphPad Prism v 10.1.0.

### Biolayer Interferometry

Sartorius HIS1K tips were submerged in 0.1% TBST for 10 minutes prior to use to hydrate tips. Tips were washed in BLI-buffer consisting of 20 mM Tris pH 7.4, 150 mM NaCl, 0.2 mM TCEP then submerged in 10 µM of 6xHis-Atg9(1-255) or 6xHis-Atg9(1-255ΔPLF) for 300 seconds to load. Tips were then washed in BLI-buffer for 60s time to establish a baseline. Tips were then submerged in Atg11NTD (1-646) at a concentration of either 10 µM, 5 µM, 2.5 µM, 1.25 µM, or 0.625 µM to allow for association for 300 seconds. Dissociation was then allowed to occur by dipping tips back into BLI-buffer for 300 seconds. The same setup was repeated with non-His loaded tips (dipping just into BLI-buffer instead of 6xHis-Atg9(1-255) or 6xHis-Atg9(1-255ΔPLFs) as a double reference control. R-eq values from BLI were plotted as averages ± standard deviation and non-linear regression fitting was done in Graphpad Prism v 10.1.0 to determine dissociation constant values.

### Cryo-EM Sample Preparation, Data Acquisition and Processing

3 µl of purified Atg11-NTD at protein concentrations of 1 mg/ml and 0.7 mg/ml with 0.05% beta-octylglucoside (OG) was applied onto one and two grids respectfully. UltrAuFoil R1.2/1.3, 300 mesh grids were glow-discharged for 1 minute at 15 mA prior to sample application. Sample application was done in a Vitrobot Mk4 set at 100% humidity, 4 ^°^C and sample preparation was done at a blot force of 4 for 3 seconds. Data was collected on a Glacios at 200 kV, 130,000x magnification, 0.89 Å pixel size with a range of −0.8 to −1.5 µm defocus. 3 to 4 images were taken per hole, depending on the dataset, with fringe-free imaging to narrow the exposed area and beam shift to move between areas in a hole. A Falcon 4i camera was set to counting mode and the images recorded as compressed tiffs. Selectris energy filter was set to 10 eV slit width. The total electron dose was 48.54 e/Å^2^.

Cryo-EM data processing was carried out using the CryoSPARC v5.0.6. An initial 8899 movies were imported into CryoSPARC and subjected to motion correction and contrast transfer function (CTF) estimation. Particles were initially picked from 500 micrographs using blob picker with a diameter of 220 to 320 Å. Particles were extracted and used for 2D classification to identify templates for template picking. 2D classes containing the Z shaped Atg11 particles were used for template picking with a diameter of 300 Å. Particles were extracted with a box size of 560 pixels and down sampled to 280 pixels for a pixel size of 1.78 Å. This initial particle set was subjected to 2D classification which revealed very limited views of Atg11. Following particle cleaning in 2D classification, an initial model was obtained via ab-initio and non-uniform refinement. This initial volume was used to generate templates for repicking the micrographs. Template picking was performed with templates generated from the initial volume and particles were then extracted with a box size of 600 pixels and down sampled to 300 pixels for a pixel size of 1.78 Å. Particles were subjected to multiple rounds of 2D classification to identify alternate views which could be combined with the primary Z shaped views. This resulted in 152,259 particles which were used for ab-initio reconstruction with three classes. Two of these classes had similar Z shaped volumes. Particles from these classes were combined into a non-uniform refinement which gave a 6.93 Å reconstruction.

A second dataset of 7936 movies was collected, imported into CryoSPARC and used for path motion correction and CTF estimation. Template picking was performed with both the original and new datasets to see if less populated alternate views could be identified during 2D classification by combining particles from datasets. 1.4 million particles and 1.2 million particles were picked from the first and second datasets, respectively. Particles were extracted with a box size of 600 pixels and down sampled to 300 pixels for a pixel size of 1.78 Å. To identify alternate views from the Z shape, particles were subjected to iterative rounds of 2D classification where the Z shaped views were removed and the remaining particles were subjected to 2D classification. Particles containing the best 2D classes for the Z shape along with all orthogonal views were then combined with the particles from the earlier reconstruction and duplicates were removed based on a separation distance of 150 Å. This left 239,340 particles which were subjected to heterogenous refinement with two classes. One of these classes, containing 111,239 particles, was used for non-uniform refinement which resulted in a 6.68 Å reconstruction. One side of the Z had much weaker density and a local refinement mask was generated focusing on one arm and the central region of the Z. The local refinement generated a 6.39 Å reconstruction which had density consistent with the helices present in the AlphaFold model of the Atg11-NTD.

### Atg11-NTD hybrid structural model generation

The AlphaFold server was used to generate an initial model of dimeric Atg11 (1-646) [61,62]. The model was opened in ChimeraX and docked into the local refinement reconstruction [63]. The map was contoured to a level of 0.4 to reveal density consistent with helical structures. Regions outside the density at a contour level of 0.3 were removed with the exception of 546-553 since clear density was visible on both sides of this region. The structure and volume were opened in Coot where we performed chain refinement to improve the overall fit of the model to the map [64]. Real space refinement was then performed using Phenix [65].

### Atg32 Clustering Assay and Vacuolar Delivery Assay

For GFP-Atg32 puncta formation experiments, *atg11Δ* cells expressing GFP-Atg32 under the MET25 promoter were grown to mid-log phase in SMD supplemented with 1 mg/mL methionine (Sigma Aldrich, M9625) [54]. GFP-Atg32 expression was then induced by shifting cells to SMD without methionine for 1 h. Subsequently, cells were subjected to nitrogen starvation (0.17% yeast nitrogen base without amino acids and ammonium sulfate; 2% glucose) for 15 minutes and imaged between 15- and 30-minutes following starvation. To quantify GFP-Atg32 puncta, a segmented line was drawn along the region of the mitochondrion displaying a potential GFP-Atg32 punctum. The intensity of the punctum was compared to the surrounding mitochondrial GFP-Atg32 background intensity, and only those puncta showing at least threefold enrichment over background were counted as bona fide puncta.

To quantify vacuolar delivery of GFP-Atg32, cells were grown similarly, starved for 90 minutes and then imaged. For analysis of vacuolar GFP, maximum-intensity projections of cells expressing GFP-Atg32 were background-corrected using the mode intensity value. Using the CMAC-stained image as a guide, one vacuole per cell was marked with a circle using the multipoint tool set to a medium circle size. Circles were placed on regions of the vacuole that did not contain mitochondria to avoid measuring mitochondrial GFP signal. GFP intensity values were normalized to the highest signal within each replicate. Data from three independent replicates were pooled and plotted as the mean, with error bars representing the standard deviation.

### OM45-GFP Processing Assay

*atg11*Δ TKYM22 cells were grown in YPD to an OD_600_ of 0.8-1 and transformed with respective plasmids via the LiAc method and plated on respective SMD plates. Plates were allowed to grow for 3-4 days [57]. Cells were then grown in 20 mL of SMD + glucose to an OD_600_ of ∼1. Cells with then shifted to grow in 50 mL of SML (0.67% yeast nitro-gen base, 2% lactic acid [Sigma Aldrich, L6661] and supplemented with the appropriate amino acid mixture) for ∼16 hours at which point 10 OD_600_ equivalents worth of cells were taken to be processed as “fed cells”. The remainder of the cells with starved in SD-N + glucose for 6 hours then 10 OD_600_ equivalents worth of cells were taken to be processed as “nitrogen starved cells”. Cells were then washed with PBS and then resuspended in 10% trichloroacetic acid and incubated on ice for 10 minutes. Cells were spun down at 18,000 g for 10 minutes at 4 ^°^C and wash 3 times with ice-cold acetone. Pellets were dried of all acetone and resuspended in 1x sample buffer and placed in tubes for bead beating for 45 seconds. Samples were then spun down at 20,000g for 5 minutes at 4 ^°^C and supernatant was recovered and boiled for 5 minutes at 95 ^°^C for Western Blotting. For GFP blots anti-GFP antibody from (Santa Cruz sc:9996) was used at 1:1000, Myc blots used and anti-Myc antibody from (abcam: 9E10) at 1:1000, and actin blots used and anti-actin antibody from (Invitrogen: MA1-744) at 1:5000. All of these were in 5% blotto in 0.1% TBST. Secondary antibody was and anti-mouse-HRP antibody from (Sigma: A4416) also in 5% blotto in 0.1% TBST at 1:20000.

## DATA AVIALIBILITY

The cryo-EM local refinement volume of the Atg11-NTD was deposited with the EMDB with ID EMD-78448.

## AUTHOR CONTRIBUTIONS

S.I.N., D.A. and M.J.R. conceived and planned the experiments. S.I.N., D.A., A.E.H., and Z.B., carried out the experiments. S.I.N, and M.J.R. drafted the manuscript. All authors edited the manuscript.

## DISCLOSURE AND COMPETING INTEREST STATEMENT

The authors declare that they have no competing interests.

## ACKNOWLEDGMENTS

This work was supported by the National Institute of General Medical Sciences (NIGMS) grant R35GM128663 to M.J.R. Purification of *E. coli* proteins was performed in part at the BioMT Molecular Tools Core at Dartmouth College which is supported by NIGMS grant P20GM113132. Sequencing of plasmids was performed by the Molecular Biology Shared Resource Center which is supported by NCI Cancer Center Support Grant P30CA023108. We thank Dr. Benedikt Westermann for the kind gift of GFP-Atg32 plasmid. We would like to thank Ann Lavanway for her work and support at the Dartmouth Imaging Facility, supported by grant NIH grant S10OD021616. We would like to thank Dr. Margie Ackerman of the Dartmouth Engineering department for help and use of Octet for Biolayer Interferometry. We would like to thank Boston University’s CryoEM Core and Dr. Chad Hicks for help and guidance in the sample preparation, grid screening and data collection of cryoEM data that was supported by NIH grant S10OD032253.

## Notes

### Competing Interest Statement

The authors have declared no competing interest.

