## Supplemental Materials for "The Z-shaped N-terminal Domain of Atg11 Coordinates Atg9 Binding and Recruitment in Selective Autophagy"

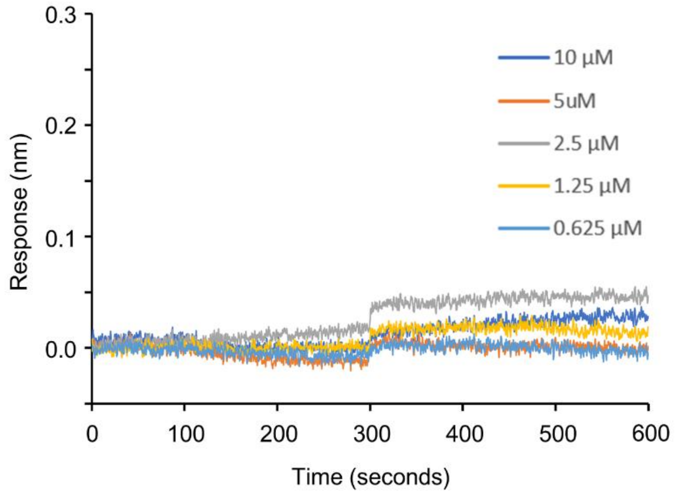


**Figure S1 – BLI traces of Atg11-NTD and 6xHis-Atg9(2-255)ΔPLF.** Representative BLI traces of 6xHis-Atg9(2-255)ΔPLFs and Atg11-NTD ranging from 0.625 µM to 10 µM.

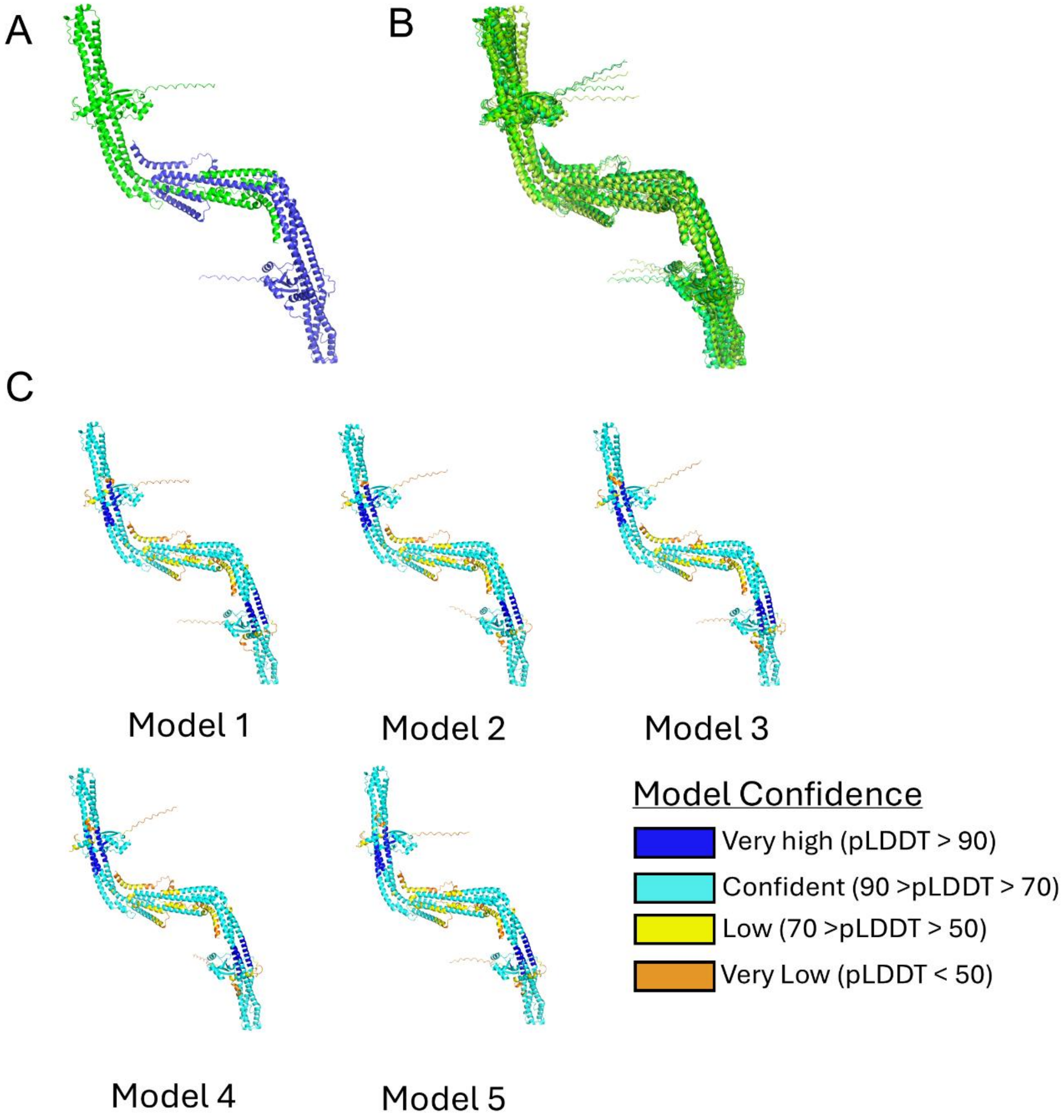


**Figure S2 – AlphaFold 3 model of the Atg11-NTD dimer.** (A) The top model is shown as a cartoon representation with one monomer in green and the second monomer in blue. (B) The overlay of the top five models of the Atg11-NTD dimer. All models are shown in different shades of green. (C) Top five models of the Atg11-NTD dimer colored based on the pLDDT confidence score where orange is very low, yellow is low, light blue is confident and dark blue is very high.

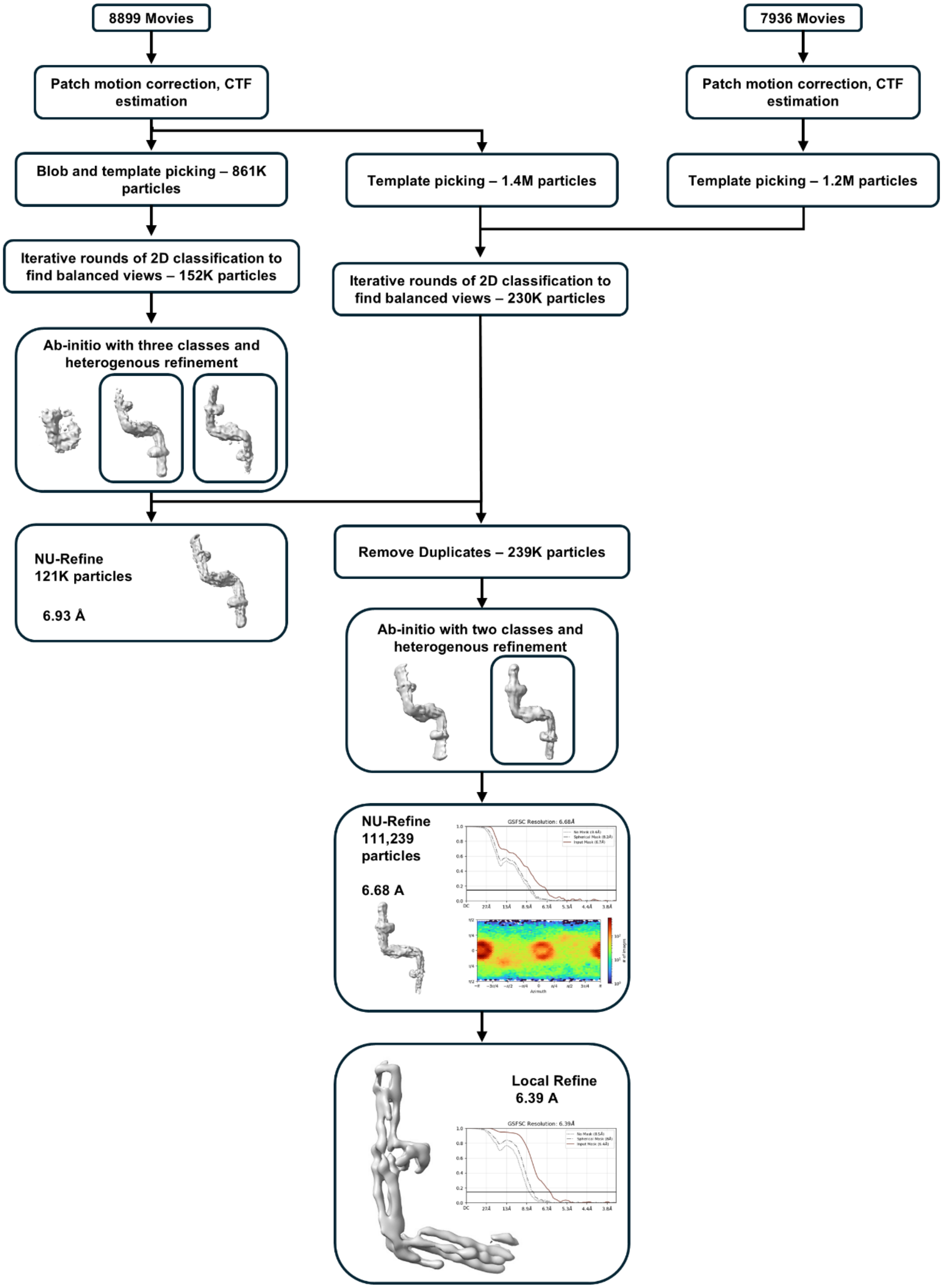


**Figure S3 – Cryo-EM processing pipeline for the Atg11-NTD.** Cryo-EM data processing was carried out using CryoSPARC. An initial dataset containing 8899 movies was subjected to motion correction and contrast transfer function (CTF) estimation. Blob picking was used to pick particles from 500 micrographs which were used for 2D classification to identify templates for particle picking. Template picking was performed on the entire dataset and particles were extracted with a box size of 560 pixels and Fourier cropped to 280 pixels. This particle set was subjected to particle cleaning in 2D classification, and an initial model was obtained via ab-initio and non-uniform refinement which showed significant orientation bias in the volume. Templates were generated from this initial volume and were used for template picking to try and identify alternate views missing from the initial rounds of picking. Particles were subjected to multiple rounds of 2D classification to identify alternate views which were then combined with the primary Z shaped views. This resulted in 152,259 particles which were used for ab-initio reconstruction with three classes. Particles from two of these classes were combined into a non-uniform refinement which gave a 6.93 Å reconstruction with reduced orientation bias. A second dataset of 7936 movies was imported into CryoSPARC and subjected to motion correction and CTF estimation. Template picking was performed with both the original and new datasets to see if less populated alternate views could be identified during 2D classification by combining particles from both datasets. 1.4 million particles from dataset 1 and 1.2 million particles from dataset 2 were picked and extracted with a box size of 600 pixels and Fourier cropped to 300 pixels for a pixel size of 1.78 Å. Particles were subjected to iterative rounds of 2D classification. After each round the primary Z shaped views were removed from the particle set and the remaining particles were subjected to 2D classification. Particles containing the best 2D classes for the Z shape along with all potential alternate views were then combined with the particles from the earlier reconstruction. Duplicate particles were removed based on a separation distance of 150 Å which resulted in 239,340 particles that were subjected to heterogenous refinement with two classes. The best class contained 111,239 particles and non-uniform refinement gave a 6.68 Å reconstruction. A local refinement mask was generated focusing on one arm and the central region of the Z. The local refinement generated a 6.39 Å reconstruction.

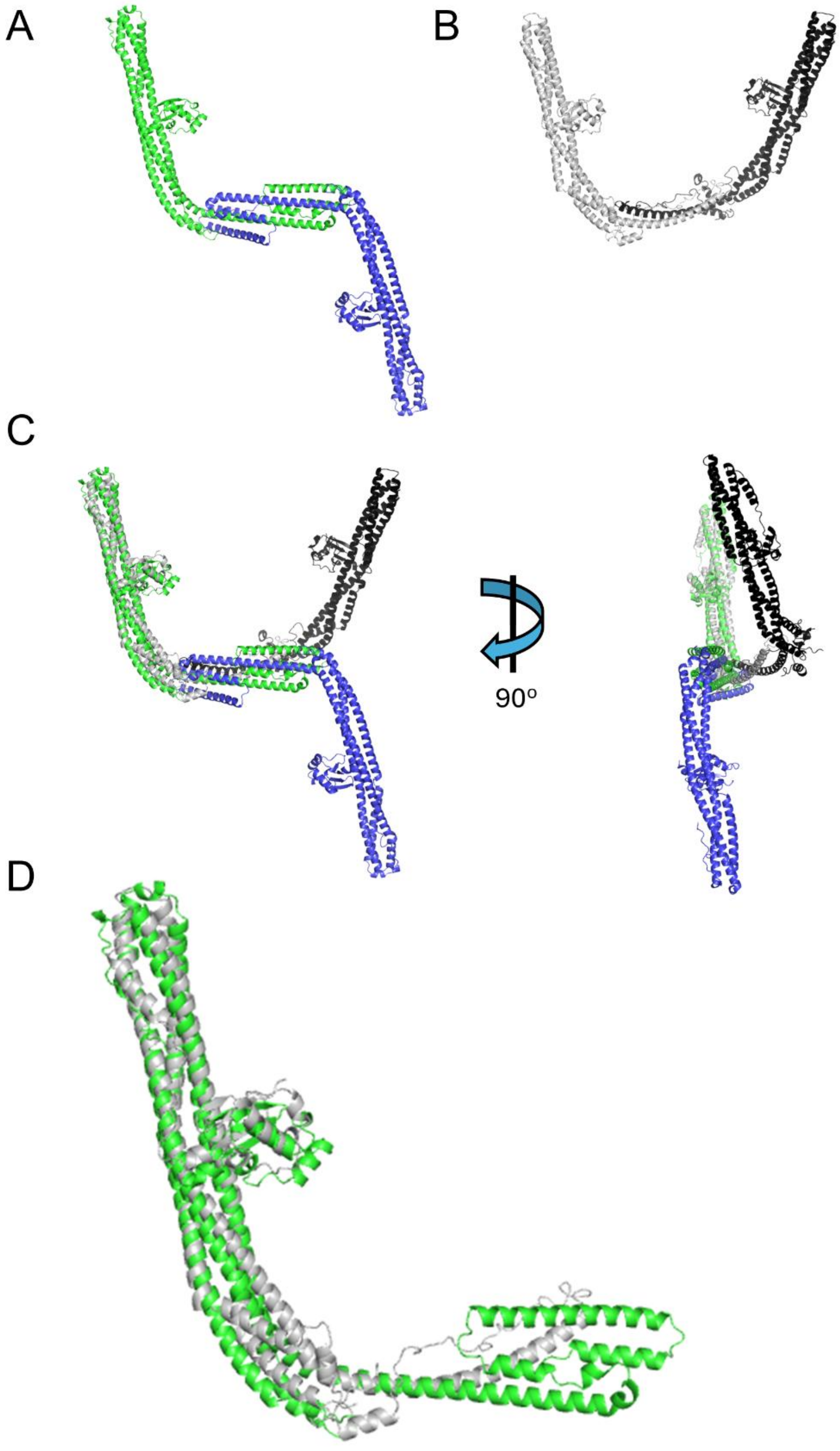


**Figure S4 – Comparison of the Atg11-NTD AlphaFold model and FIP200 structure.** (A) The top AlphaFold 3 model of Atg11-NTD is shown with all regions removed that were not visible in the density. One monomer is colored green and the second monomer is colored blue. (B) The FIP200-NTD dimer structure from PDB: 8SOI shown as a cartoon representation with one monomer colored in grey and the other colored in black. (C) An overlay of the Atg11-NTD AlphaFold model and the FIP200-NTD structure from A and B. Two perpendicular views are shown to highlight the structural differences between the dimers. (D) Overlay of the hybrid structural model of the Atg11-NTD (green) from Figure 3 with the monomer of FIP200-NTD from PDB: 8SOI (grey).

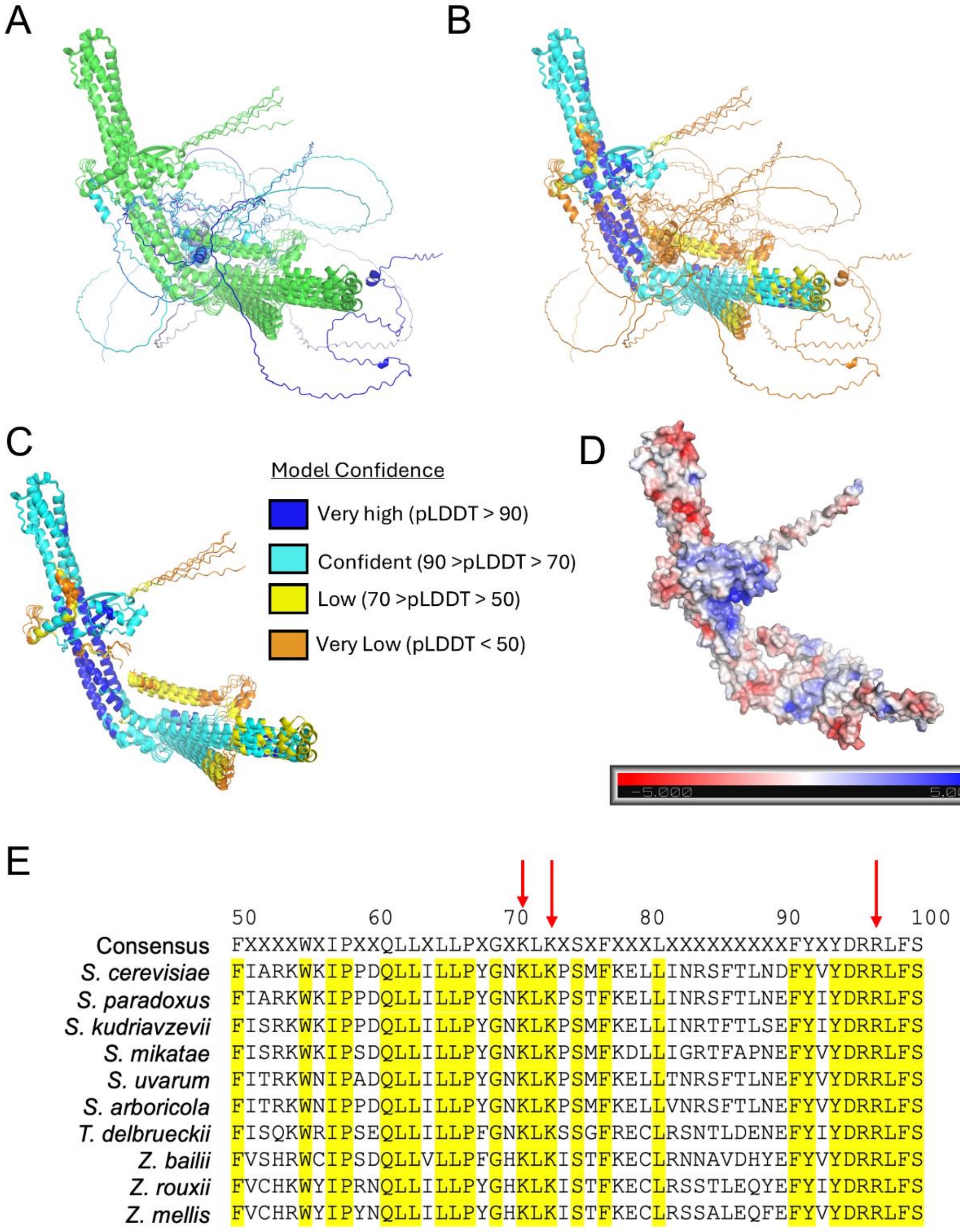


**Figure S5 – AlphaFold 3 model of the Atg9(2-255) and Atg11-NTD complex.** (A) The top five models of the Atg11-NTD and Atg9(2-255) complex are shown as an overlay with the monomer of Atg11-NTD that is not binding to Atg9 not shown. Atg11-NTD is shown in different shades of green and Atg9 is shown in different shades of blue. (B) The same overlay shown as in A except that everything is colored based on the pLDDT confidence score where orange is very low, yellow is low, light blue is confident and dark blue is very high. (C) The same representation as B except that the amino acids of Atg9(2-255) that are not binding to Atg11-NTD are hidden. (D) Electrostatic surface representation of the top model of the Atg11-NTD from C where blue is positively charged, red is negatively charged and grey is not charged. (E) Sequence alignment of ten different yeast Atg11 sequences. Amino acids that are fully conserved are highlighted in yellow. The three amino acids mutated in the CR mutant are noted with red arrows.

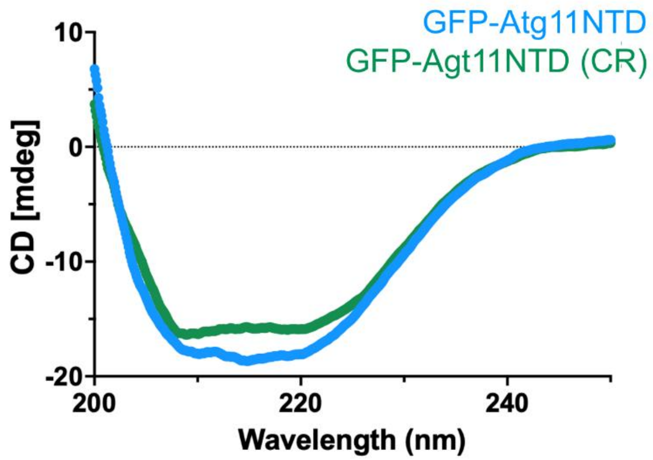


**Figure S6 – Circular dichroism of GFP-Atg11-NTD and GFP-Atg11-NTD(CR).** Circular dichroism plots are shown from 200 to 250 nm for the GFP-Atg11-NTD in blue and GFP-Atg11-NTD (CR) in green.

### Supplementary Tables

**Table S1 – Cryo-EM data collection statistics for the Atg11-NTD**.

| Atg11-NTD (Local Refinement) | |
|---|---|
| Data collection | |
| Microscope | Glacios II |
| Detector | Falcon 4i (counting mode) |
| Nominal Magnification | 130,000 |
| Voltage (kV) | 200 |
| Total Exposure ($e^-/Å^2$) | 48.54 |
| Number of frames | 35 |
| Selectris Energy filter width (eV) | 10 |
| Defocus range (µm) | -0.8 to -1.5 |
| Calibrated physical pixel size (Å/px) | 0.89 |
| Reconstruction and Refinement | |
| Particles | 111,239 |
| Resolution (Å) | 6.31 |
| Data Deposition | |
| EMDB code | 78448 |

**Table S2 – Plasmids used in this study**.

| Plasmid Name | Description | Citation |
|---|---|---|
| pSN060 | pET21 Atg11NTD(1-646)-TEV-6xHis | This Study |
| pSN040 | pET His6 TEV LIC cloning vector (1B) 6xHis-Atg9(2-255)-TEV-MBP | This Study |
| pSN054 | pET His6 TEV LIC cloning vector (1B) 6xHis-Atg9(2-255(ΔPLF1(163-165) and ΔPLF2(187-189))-TEV-MBP | This Study |
| pSN074 | pET His6 GFP TEV LIC cloning vector (1GFP) Atg11-NTD(1-646-CR (K71E, K73E, R96E)) | This Study |
| pSN021 | pET His6 GFP TEV LIC cloning vector (1GFP) 6xHis-GFP-TEV-Atg11NTD (1-646) | This Study |
| pSK073 | pET His6 TEV LIC cloning vector (1B) 6xHis-GFP-TEV-Atg11CTR (698-1178) | (Andhare et al., 2026) |
| pZB051 | pET-3a TS-mScar-Atg9(1-255)-trifold-6xHis | This Study |
| pZB036 | pGEX-4T-1 10xHis-GST-TEV-Atg9(1-255) | (Bekkhozhin, Leary, & Ragusa, 2026) |
| pZB078 | yCPlac111-Atg9-mScarlet | This Study |
| pSN007 | yCPlac33-2xGFP-Atg11 | This Study |
| pSN008 | yCPlac33 -2xGFP-Atg11-NTD(2-646) | This Study |
| pSN010 | yCPlac33-2xGFP-Atg11-CTR (699-1178) | This Study |
| pSN019 | yCPlac33-2xGFP | This Study |
| pHMP001 | yCPlac33-2xGFP-Atg11-NTD(2-646)-µNS | This Study |
| pZG001 | yCPlac111-3xMyc-Atg11 | This Study |
| pSN069 | yCPlac33-2xGFP-Atg11- CR(K71E, K73E, R96E) | This Study |
| pSN080 | yCPlac111-3xMyc-Atg11-CR(K71E, K73E, R96E) | This Study |
| GFP-Atg32 | pRS415-MET25-GFP-Atg32 | (Böckler & Westermann, 2014) |
| pDA078 | pCu415-3xFLAG-mScarlet | This Study |
| pKL21 | pCu415-Atg9-HA | This Study |

**Table S3 – Yeast strains used in this study**.

| **Strain Name** | **Genotype** | **Citation** |
|---|---|---|
| BY4742 *atg11*Δ | BY4742 MATα his3Δ1 leu2Δ0 lys2Δ0 ura3Δ0 *atg11*Δ::Kan | Yeast KO Collection (Thermo Fisher Scientific) |
| SNY009 | BY4742 MATα his3Δ1 leu2Δ0 lys2Δ0 ura3Δ0 *atg9*Δ::Kan, *atg11*Δ::His | This Study |
| SNY019 | BY4742 MATα his3Δ1 leu2Δ0 lys2Δ0 ura3Δ0 *atg11*Δ::Kan, *atg9*Δ::His, *atg17*Δ::Nat | This Study |
| XXY003 | SEY6210 (MATα leu2-3,112 ura3-52 his3-Δ200 trp1-Δ901 suc2-Δ9 lys2-801; GAL) OM45-GFP::TRP1, *atg11*Δ::Nat | (Margolis, Katzenell, Leary, & Ragusa, 2020) |
